# Respiratory Syncytial Virus (RSV) optimizes the translational landscape during infection

**DOI:** 10.1101/2024.08.02.606199

**Authors:** Kyra Kerkhofs, Nicholas R. Guydosh, Mark A. Bayfield

## Abstract

Viral infection often triggers eukaryotic initiator factor 2α (eIF2α) phosphorylation, leading to global 5’-cap-dependent translation inhibition. RSV encodes messenger RNAs (mRNAs) mimicking 5’-cap structures of host mRNAs and thus inhibition of cap-dependent translation initiation would likely also reduce viral translation. We confirmed that RSV limits widespread translation initiation inhibition and unexpectedly found that the fraction of ribosomes within polysomes increases during infection, indicating higher ribosome loading on mRNAs during infection. We found that AU-rich host transcripts that are less efficiently translated under normal conditions become more efficient at recruiting ribosomes, similar to RSV transcripts. Viral transcripts are transcribed in cytoplasmic inclusion bodies, where the viral AU-rich binding protein M2-1 has been shown to bind viral transcripts and shuttle them into the cytoplasm. We further demonstrated that M2-1 is found on polysomes, and that M2-1 might deliver host AU-rich transcripts for translation.

**Importance:** Viruses strongly rely on the host’s translational machinery to produce viral proteins required for replication. However, it is unknown how viruses that do not globally inhibit cap-dependent translation compete with abundant host transcripts for ribosomes. In this study, we found that respiratory syncytial virus (RSV) infection results in redistribution of 80S monosomes into the polysomes. High-throughput sequencing of translating transcripts revealed that low translation efficiency transcripts become more efficient at ribosome recruitment which are virus-resembling AU-rich host transcripts. Finally, we also uncover that AU-rich RNA binding protein RSV-M2-1 interacts with polysomes through contacts to mRNA. These findings revealed that RSV optimizes the translational landscape rather than inhibiting host translation.

## Introduction

Viral infection often results in remodeling of the host’s translational landscape caused by viral proteins that hijack translation regulatory factors, presence of high numbers of viral transcripts and host-induced innate immune activation. Viruses rely completely on the host’s ribosomes for viral protein translation and thus compete with host mRNAs ^1^. Cells exposed to stress often regulate gene expression through 5’-cap-dependent translation arrest, often mediated through phosphorylation of the α-subunit of eIF2 (eIF2α) by stress-activated kinases leading to inhibition of subsequent rounds of initiation. Without translation initiation, ribosome-free transcripts are bound by RNA-binding proteins and assemble into stress granules ^2,3^.

Respiratory syncytial virus (RSV) is an enveloped virus containing a non-segmented, single-stranded, negative-sense RNA genome expressing 10 individually 5’-capped and polyadenylated transcripts transcribed by the viral polymerase ^4–7^. Following fusion of the viral particle with the host’s membrane, the nucleocapsid is released into the cytoplasm and the viral polymerase (containing phosphoprotein RSV-P and large polymerase protein RSV-L) starts replication and transcription of the viral genome ^8,9^ within cytoplasmic membraneless inclusion bodies ^10–12^. Transcription of RSV transcripts requires an additional protein, RSV-M2-1, which functions as a transcription processivity factor ^13,14^. M2-1 has also been shown to bind nascent transcribed viral transcripts and transport these from inclusion bodies into the cytoplasm ^12,15,16^. Translation of the viral transcripts occurs in the cytoplasm using the host’s ribosomes^12,17^.

Since RSV transcripts mimic post-transcriptional features of host transcripts ^18^, it would be detrimental to viral gene expression if 5’-cap-dependent translation initiation were inhibited through eIF2α phosphorylation by stress-activated kinases. RSV infection results in both upregulation of the stress-activated kinase PKR ^19–21^ and PKR activation through dimerization and autophosphorylation ^22,23^. This normally induces eIF2α phosphorylation leading to reduced translation initiation and stress granules formation. Although multiple studies have demonstrated that RSV has developed different strategies to maintain host translation levels by negating eIF2α phosphorylation ^20,24–26^, another study reports that RSV induces stress granules ^27^. Despite elucidation of inhibitory eIF2α phosphorylation strategies, stress granule formation during RSV infection remains controversial. Additionally, since RSV does not induce a strong “host shutoff” inhibiting global 5’-cap-dependent translation initiation ^20,28^, it remains to be determined how RSV successfully competes with host transcripts for the machinery required for translation of its viral genes.

In this study, we describe host translatome changes during infection towards preferential translation of transcripts more similar to viral transcripts. We first confirm that RSV limits inhibition of widespread translation initiation seen by lack of both eIF2α phosphorylation and stress granule formation. Interestingly, we found that the number of ribosomes within polysomes increases during infection, indicating enhanced ribosome loading. Next, through high-throughput sequencing of total and polysome-associated transcripts we describe how transcripts that are normally lowly translated become more efficient at recruiting ribosomes during infection. We show that more efficiently translated host transcripts are AU-rich, similar to viral transcripts. In addition, we found that AU-binding protein RSV-M2-1 is present on polysomes, and that M2-1 might also deliver host AU-rich transcripts for translation.

## Results

### Ribosome occupancy is increased during RSV infection

While RSV activates the stress-induced eIF2α-phosphorylating kinase PKR ^20,21^, the extent of downstream phosphorylation of eIF2α and consequent stress granule formation remains unclear ^20,21,24–27^. We tested if eIF2α is phosphorylated during infection and found that only a small fraction is phosphorylated by western blot (**Figures 1A** and **S1A**) compared to cells treated with arsenite (NaAsO_2_) (**Figures 1B** and **S1B**). Similar results were observed at earlier RSV-infection timepoints (**Figure S1C**). To further validate that eIF2α remains unphosphorylated during infection, we also confirmed the absence of stress granules in RSV-infected cells by indirect immunofluorescent staining using stress granule markers PABP and G3BP (**Figures S1D-E**) ^29^, which is in stark contrast with NaAsO_2_-treated cells (**Figure S1F**). Consistent with previous work ^12^, we observed that following RSV infection, cytoplasmic inclusion bodies were formed which function as sites of viral RNA transcription by the viral polymerase and consists of viral proteins N, P, L and M2-1 and a selection of host proteins with functions in translation, including PABP (**Figure S1D**, zoom), but excluding *bona fide* stress granule marker G3BP (see **Figure S1E**, zoom) ^12,24^.

**Figure 1.**
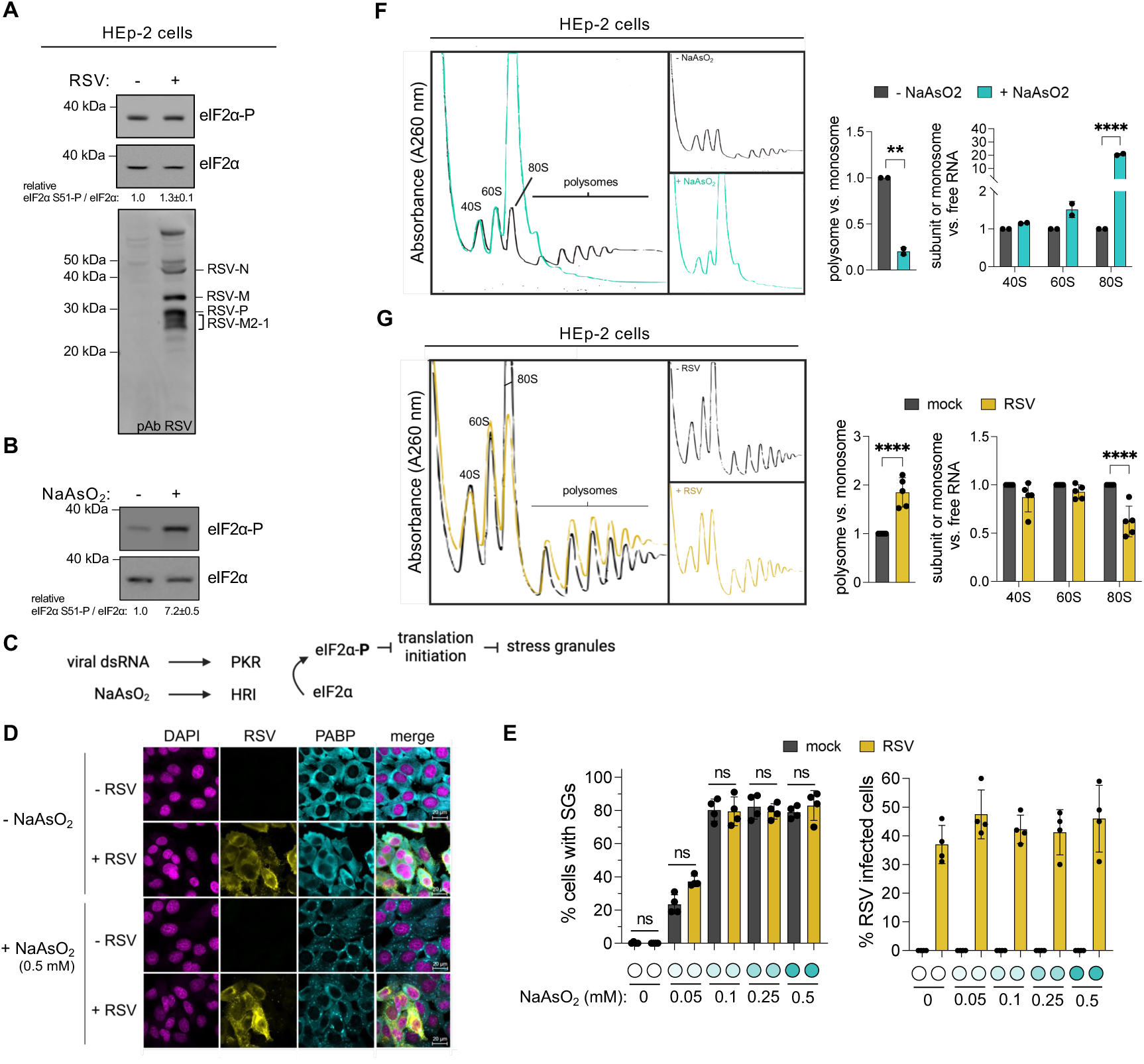
RSV infection maintains translation and increases ribosome occupancy. **(A,B)** Western blot comparing eIF2α-P and total eIF2α levels between (**A**) mock-and RSV-infected (MOI 1, 24h) and (**B**) untreated and NaAsO_2_-treated (positive control) (0.5 mM, 1h) HEp-2 cells. Relative quantification against control cells is shown below (n = 3). RSV infection was confirmed by immunoblotting with a polyclonal anti-RSV antibody (pAb). **(C)** Schematic representation of eIF2α-phosphorylating kinases activated during NaAsO_2_ stress (HRI) and viral infection (PKR). **(D,E)** RSV does not inhibit NaAsO_2_-induced stress granule formation. Indirect immunofluorescent staining of mock-and RSV-infected (MOI 1, 24h) and NaAsO_2_-treated (0.5 mM, 1h) HEp-2 cells (n = 3) (**D**). DAPI staining identifies nuclei, PABP detects stress granules and RSV infected cells were detected with a polyclonal RSV antibody. (**E**) Quantification of mock-and RSV-infected HEp-2 cells (MOI 1, 24h) treated with different concentrations of NaAsO_2_ (1h). More than 200 cells were quantified at 20X magnification (n = 2). Number of cells were determined by DAPI, stress granules by PABP and RSV infection by polyclonal RSV staining. P values were calculated with one-way ANOVA with Tukey’s multiple comparisons test. **(F,G)** RSV redistributes 80S monosomes to polysomes. Polysome profiles of (**F**) untreated and NaAsO_2_-treated (0.5 mM, 1h) and (**G**) mock-and RSV-infected (MOI 1, 24h) sucrose gradient fractionated HEp-2 cells. AUC quantification between polysomes and monosomes (40S, 60S and 80S) are plotted to estimate translation levels. AUC quantification between free RNA fraction (not shown) and 40S, 60S and 80S are plotted to determine changes in free monosomes and 80S subunits. AUC: area under the curve. P values were calculated with an unpaired t-test for polysome vs monosome comparisons and a two-way ANOVA with Šídák’s multiple comparisons test. See also **Figure S1.**

To further test if lack of stress granule formation by RSV is caused by inhibition of eIF2α phosphorylation or by rapid dephosphorylation of eIF2α-P, we used NaAsO_2_ to activate another eIF2α-phosphorylating kinase, HRI (as opposed to PKR which recognizes viral dsRNA), leading to eIF2α phosphorylation, reduced translation initiation and stress granule formation (**Figure 1C**) ^30^. We found that RSV-infected cells retained the ability to form stress granules after NaAsO_2_ treatment (**Figure 1D**), consistent with previous work ^17^. Next, the same experiment was performed with lower NaAsO_2_-concentrations to ensure that activation of the NaAsO_2_-activated stress signalling pathways was not overwhelming any potential RSV-induced inhibitory system. Consistent with the highest NaAsO_2_ concentration, we found no significant differences in stress granule formation between mock-and RSV-infected cells after NaAsO_2_ treatment (**Figure 1E**), suggesting that infected cells are capable of stress granule formation but without inducing them during RSV infection.

Next, we performed polysome profiling to separate mRNAs according to the number of bound ribosomes. By fractionating lysates on sucrose gradients, we obtained separation between free RNA (not shown), 40S and 60S ribosomal subunits, 80S monosomes, and polysomes (**Figures 1F-G**). Treatment with NaAsO_2_ results in a strong translational arrest seen by a large increase in the 80S peak and disappearance of polysomes (**Figure 1F**), consistent with translation inhibition as shown previously ^31,32^. We expected to observe similar levels in polysomes between mock-and RSV-infected lysates, but interestingly we found that polysome levels are consistently increased as seen by an increase in the polysome/monosome ratio compared to mock-infected cells across all replicates (**Figures 1G** and **S1G**). The increase in polysomes is accompanied by a decrease in 80S monosomes, while 40S and 60S subunit levels remain similar (**Figures 1G** and **S1G**), indicating that 80S monosomes are being redistributed to the polysomes as opposed to an increased level of ribosome production. Overall, our findings demonstrate that during RSV infection, stress granules are absent, and ribosome occupancy is increased.

### RSV infection induces three distinct modes of host translation changes

To determine which transcripts are associated with polysomes during infection we isolated total and polysome-associated mRNA from mock-and RSV-infected cells and performed high-throughput sequencing after poly(A) (A+) enrichment (**Figures 2A** and **S2**, **Table S1**). Next, we determined the relative abundance of DESeq2 normalized reads (**Tables S2** and **S3**) ^33^. After plotting normalized reads by transcript type for mock-and RSV-infected cells, we observed that viral transcripts occupy approximately 14% of total A+ RNA and 1.5% of polysomal A+ RNA at 24 hours post-infection (**Figure 2B**). Next, we determined the expression levels of viral transcripts in comparison with host transcripts by plotting the distribution of normalized reads of all 10 viral transcripts and each individual protein-coding mRNA for mock-and RSV-infected samples (excluding viral mRNAs) (**Figure 2C**). In total A+ mRNA samples, most viral transcripts are present at higher abundance than the highest expressed host mRNA (**Figure 2C**, left). We next investigated whether host and viral transcripts were translated with similar efficiency and found that viral transcripts in the polysomal A+ mRNA fractions were found at levels similar to highly expressed host transcripts (**Figure 2C**, right). In conclusion, viral transcripts are found within polysomes to the same extent as highly translated host transcripts, indicating that a high number of viral proteins are being produced. However, viral transcripts appear to not be as efficient at recruiting ribosomes, as seen by a large difference between their abundance in total and polysomal A+ RNA fractions (see later).

**Figure 2.**
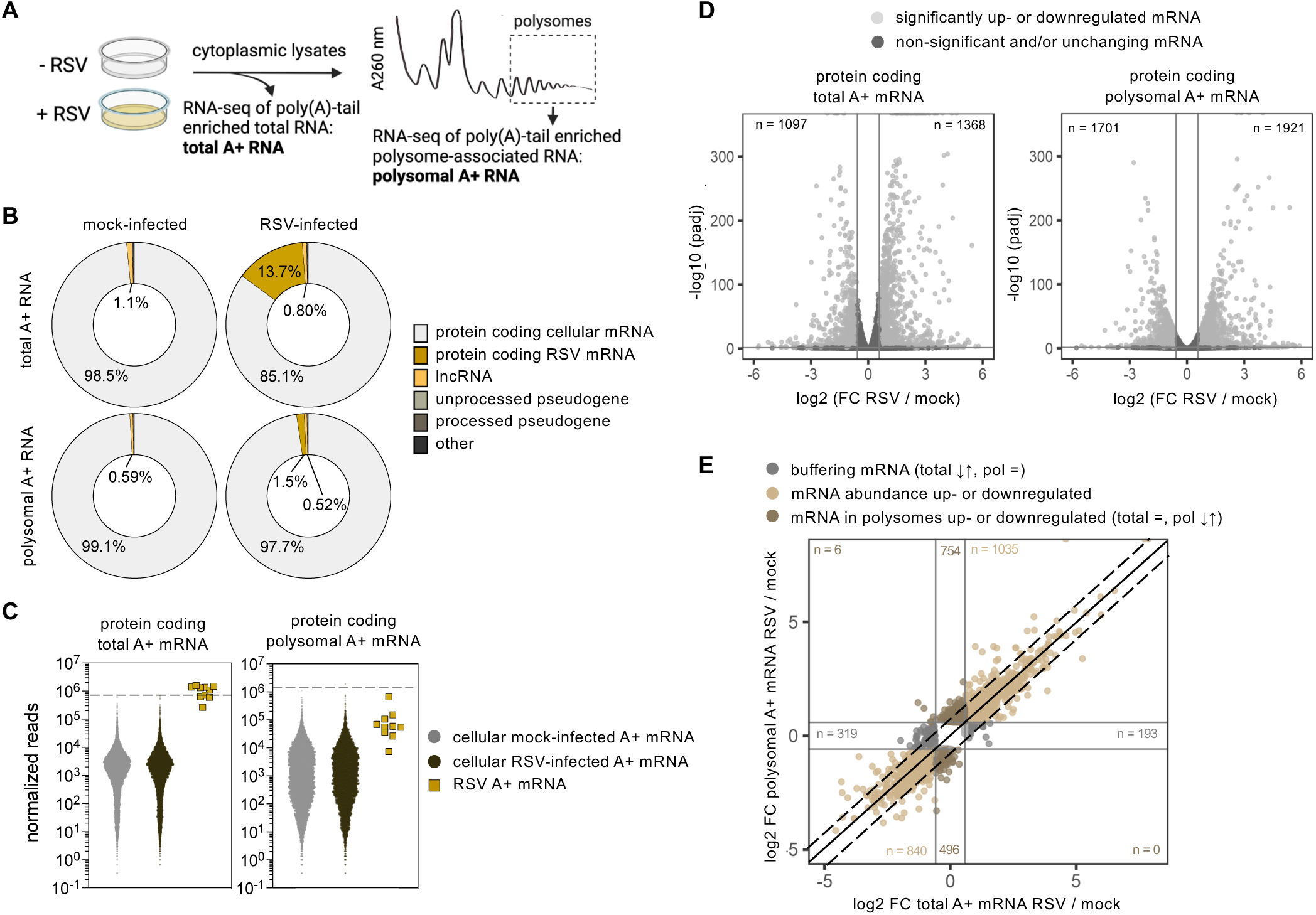
RSV infection induces three distinct modes of host translation changes. **(A)** Schematic representation of experimental design. Cells were mock- (-RSV) or RSV-infected (+ RSV) with a multiplicity of infection (MOI) of 1 (*i.e.* one viral particle per cell) for 24h. Prior to harvest, cells were treated with cycloheximide (CHX) to halt translation elongation and stabilize ribosomes on mRNA. Cell lysates were fractionated on sucrose gradients separating 40S, 60S, 80S and polysomes. RNA was isolated from heavy polysomes and from total RNA (acquired prior to fractionation), poly(A)-tail enriched (A+) and analyzed by next-generation sequencing. **(B)** Pie chart demonstrating the distribution of different RNA types (averaged biological triplicates) after DESeq2 normalization. A+: poly(A)-tail enriched RNA. **(C)** Distribution of DESeq2 normalized protein-coding reads (averaged biological triplicates) for mock-and RSV-infected cells for total (*left*) and polysomal A+ mRNA (*right*). The dotted line indicates the highest expressed host transcript in mock-infected cells. **(D)** Volcano plots of differentially expressed protein-coding host mRNAs comparing mock-and RSV-infected samples (MOI 1, 24h) (three biological replicates) from total A+ mRNA (*left*) and polysomal A+ mRNA (*right*). The horizontal line indicates a cutoff of padj < 0.05 and vertical lines indicate a 1.5-FC. FC: fold change. **(E)** Scatterplot between the log2 FC of total mRNAs (RSV / mock) and polysome associated mRNAs (RSV / mock). Horizontal and vertical lines indicate a 1.5-FC. Diagonal lines indicate transcripts with changing translation efficiencies (TE) (see later). FC: fold change. See also **Figure S2.**

Next, to determine how total and polysomal transcripts are affected by RSV infection, we performed differential expression analysis between RSV-and mock-infected samples using DESEq2 for both total and polysomal A+ RNA (see methods). We determined differentially expressed protein-coding transcripts (padj < 0.05 and fold-change (FC) < or > 1.5-fold) and plotted these on volcano plots (**Figure 2D**). We observed that many host transcripts (both total and polysomal) are significantly up-or downregulated during infection (**Figure 2D**). Although it is well known that RSV infection induces multiple host responses that activate or repress transcription of host genes ^34^, resulting in differentially expressed genes between mock-and RSV-infected cells (**Figure 2D**, left), how polysome-associated transcripts change during RSV infection remains to be investigated. The number of polysome-associated transcripts determines the amount of protein produced, which makes this a critical gene regulatory step for the cell.

Changes in polysome-associated mRNA abundance can be caused by two factors. First, changes in total transcript abundance tend to cause corresponding changes in polysome association. Second, through enhanced (or decreased) ribosome recruitment, independent of total mRNA fluctuations, transcripts will also be increased (or decreased) in polysomes. To further understand how transcripts are being enriched or depleted within polysomes during infection, we plotted the fold change (FC) of differentially expressed transcripts (padj < 0.05) between RSV-and mock-infected samples of polysomal A+ mRNAs against total A+ mRNAs (**Figure 2E**). We found that most of the changes in polysomal mRNA abundance were driven by changes in total A+mRNA, seen by the distribution of datapoints along the diagonal (**Figure 2E**, mRNA abundance up-or downregulated, light brown). In addition, a high number of transcripts changed in their polysome association with no or opposite changes in total abundance (**Figure 2E**, mRNA in polysomes up-or downregulated, dark brown). And lastly, we also found some transcripts that changed in total mRNA abundance but without corresponding changes in their association with polysomes, termed translational buffering (**Figure 2E**, buffering, dark grey). During buffering, changes in translation compensate for changes in total mRNA abundance. As a result, any RSV-induced abundancy changes of these transcripts are buffered at the translational level and thus will not result in changes in protein production. Overall, these data indicate that during RSV infection all three major types of translational regulation mechanisms occur.

### Transcripts with low TE become more efficient at recruiting ribosomes than those with high TE during RSV infection

Given our observations above that some mRNAs in the total pool were selectively enriched or depleted in the polysomes, we quantified the translatability of host protein-coding mRNAs with the translation efficiency (TE) metric in mock-and RSV-infected cells. The TE is calculated by taking ratio between polysomal and total A+ mRNAs and is a measure of how well mRNAs become loaded with ribosomes (**Figure 3A**, *top*, **Table S4**). For example, in **Figure 2E**, transcripts with a substantially increased TE during infection are found above the upper dashed diagonal line and those with substantially lower TE are found under the lower diagonal line. We plotted the TE of all protein-coding transcripts from RSV-against mock-infected samples (**Figure 3A**, scatterplot; replicates shown in **Figure S3A**) and computed a histogram of the ratio of these values at each data point (**Figure 3A**, bar chart). Intriguingly, the data did not fall stochastically around the diagonal of the scatterplot but exhibited a clear pattern where the points generally fell above the diagonal for low TE transcripts and below for high TE transcripts. To better quantify this observation, we divided the plot based on a high (> 2) and low TE (< 2) in uninfected cells (**Figure 3B**), where, for example, a transcript considered to have a high TE is enriched at least two-fold in polysomes compared to the total found in the cell. More specifically, transcripts that are heavily translated in uninfected cells are likely more efficient at recruiting ribosomes and thus obtain a high TE (> 2) while transcripts that are less efficient at recruiting ribosomes obtain a low TE (< 2). This division further demonstrates that during RSV infection, normally highly translated host transcripts specifically appear to be less efficient at recruiting ribosomes (**Figure 3B**, high TE), seen as a downwards curve from the diagonal when comparing the TE between mock-and RSV-infected cells. This decrease is also reflected in the histogram below the scatterplot (**Figure 3B**). While transcripts with a high TE undergo a strong decrease in TE during viral infection, the opposite trend is observed for transcripts with a low TE (**Figure 3B**, low TE). Overall, we observe more efficient recruitment of polysomes to many low TE transcripts (n = 7007) and a relative decrease in TE of high TE transcripts (n = 3958) during RSV infection. We validated our method for assaying TE, by comparing GC% and transcript length in our data. Generally, transcripts with a higher GC content ^35^ and shorter coding sequence (CDS) lengths ^36,37^ have a higher translatability. We plotted these features for the low and high TE datasets and confirm that the high TE transcripts (>2) contain significantly shorter CDSs and higher GC-content (**Figure S3B**).

**Figure 3.**
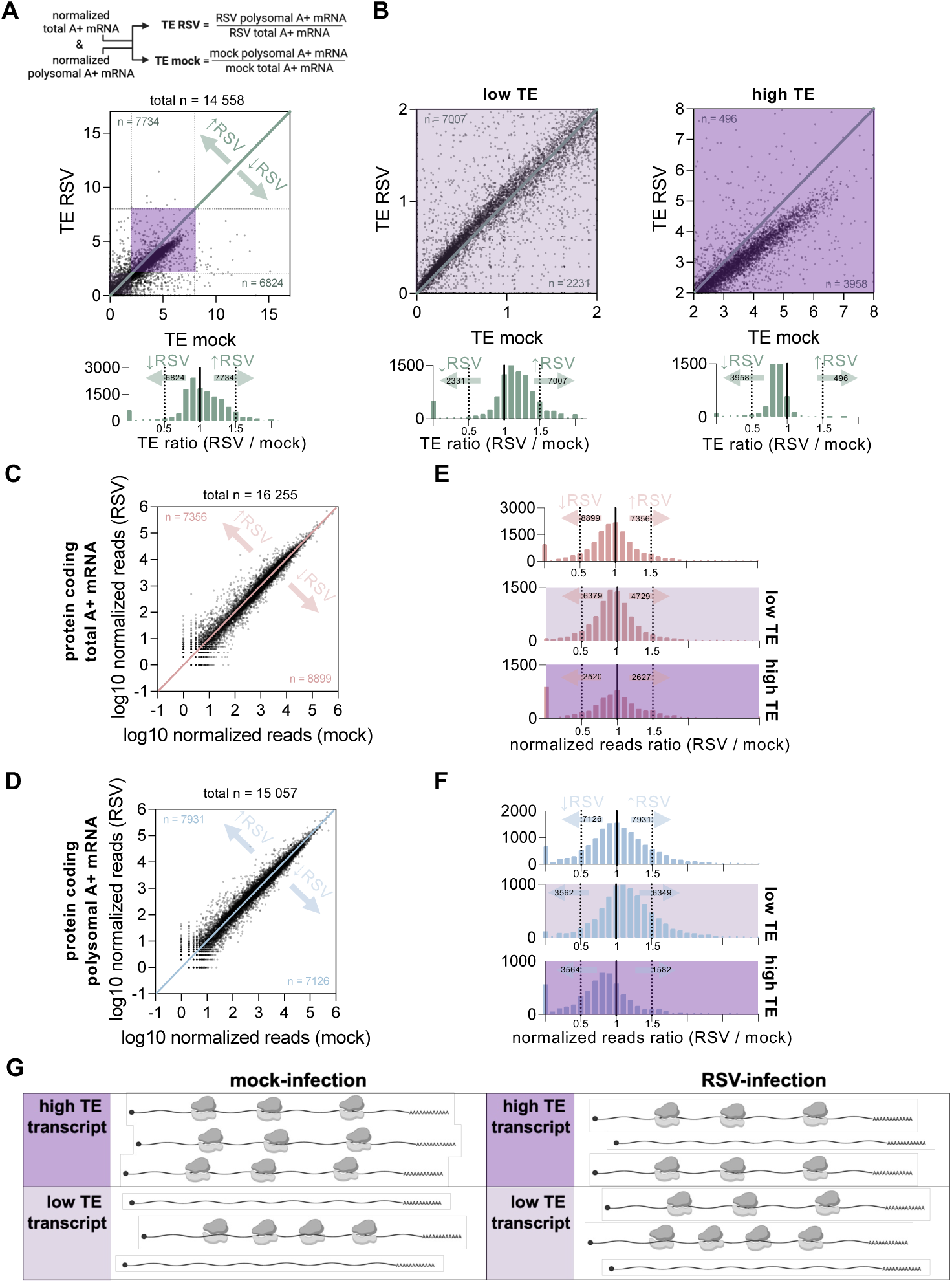
Transcripts with low TE become more efficient at ribosome recruitment during RSV infection. **(A,B)** Schematic of TE calculation. DESeq2 normalized reads obtained from experiment in Figure 2A were used to calculate ratios (**A**, *top)*. Scatterplots of TE between mock-and RSV-infected samples with a global overview (**A***, bottom)* and zoomed versions (**B**). Corresponding histograms are shown below each graph representing the fold-change between TE for mock-and RSV-infected samples. TE: translation efficiency. **(C,D)** Scatterplots of normalized reads for total cytoplasmic mRNAs (**C**) and polysome-associated mRNAs (**D**) between mock-and RSV-infected samples (MOI 1, 24h) showing even distribution along the diagonal. **(E,F)** Histograms displaying the distribution of normalized reads between mock-and RSV-infected samples for all, low TE (< 2) and high TE (> 2) transcripts for total **(E**) and polysomal A+ mRNA (**F**). This shows that changes in TE are driven by changes abundance in polysome-associated transcripts. **(G)** Schematic representation summarizing results from **A-F**. See also **Figure S3.**

Since changes in TE are driven by changes in either polysomal or total A+ mRNA (or both), we investigated relative changes in total (**Figure 3C**) and polysomal A+ mRNA (**Figure 3D**) in RSV-against mock-infected cells. Since the normalization for these plots included viral mRNAs, downward shifts of host mRNA levels tend to reflect changes in the relative proportion of reads mapping to viral RNAs in infected cells. These shifts are small for both total and polysomal A+ mRNAs since the relative proportion of viral mRNAs is limited (<14% and <2%, respectively, see **Figure 2B**). This is quantified in a histogram of RSV/mock ratio for each mRNA (**Figures 3E-F**, *top*). We then quantified the shifts for the subsets of high and low TE transcripts. We found that changes in the abundance (total A+ mRNAs) were independent of TE changes, while the changes in the polysomal A+ mRNAs correlated with changes in TE (**Figures 3E-F**). This shift is consistent with our observation above that low TE transcripts generally increase in polysome recruitment while high TE transcripts generally decrease in polysome recruitment upon infection (see **Figures 3A-B**), and that there is no simultaneous change in total mRNA levels to buffer this effect. Overall, this indicates that host transcripts with a high TE that are normally highly capable of recruiting ribosomes become less efficient—relative to low TE transcripts—at getting translated during RSV infection (**Figure 3G**). As this effect is relative, it is possible that it is driven by high TE transcripts reducing ribosome loading, low TE transcripts increasing ribosome loading, or a combination of both.

### VSV also induces a redistribution of ribosomes towards transcripts with low TE despite global “host shut-off”

Like RSV, vesicular stomatitis virus (VSV) transcribes monocistronic 5’-capped and polyadenylated transcripts ^38^. In addition, transcript features such as GC% and length within the 5’-UTR, CDS and 3’-UTR are very similar between both viruses (**Figure 4A**). Previous studies demonstrated that VSV infection induces “host shutoff” resulting in a global reduction in host mRNA abundance ^39–43^ and through inhibition of host mRNA translation, without affecting viral translation ^44,45^. Overall, as expected, VSV infection results in a reduced efficiency of host mRNAs at recruiting ribosomes ^46^ driven by a redistribution of host ribosomes onto viral mRNA ^47^. While a major ribosome redistribution occurs from host to viral transcripts, we further investigated a previously published high-throughput sequencing dataset of VSV infected total and polysomal mRNAs ^47^ to identify any ribosome redistribution trends within host mRNAs. First, we calculated the TE for mock-and VSV-infected cells, as done in **Figure 3A**. We found that the changes in TE for VSV followed a similar trend compared to RSV (**Figure 4B**). Similar to RSV, transcripts with a high TE (> 1.5) (TE cut-off determined in **Figure S4A**) appear to be the least efficient at recruiting ribosomes (**Figure 4B**, strong distribution towards the left in the histogram), compared to transcripts with a low TE (< 1.5) (**Figure 4B**, mild distribution towards the left in the histogram). We note that most host transcripts in VSV-infected cells are lower in TE compared to mock-infected cells, seen as datapoints mostly below the diagonal (**Figure 4B**, n = 7806 downregulated, n = 1843 upregulated). This likely reflects the global inhibition of host mRNA translation initiation in favor of viral initiation, even as trends for high and low TE mRNAs are otherwise similar to RSV, as noted above.

**Figure 4.**
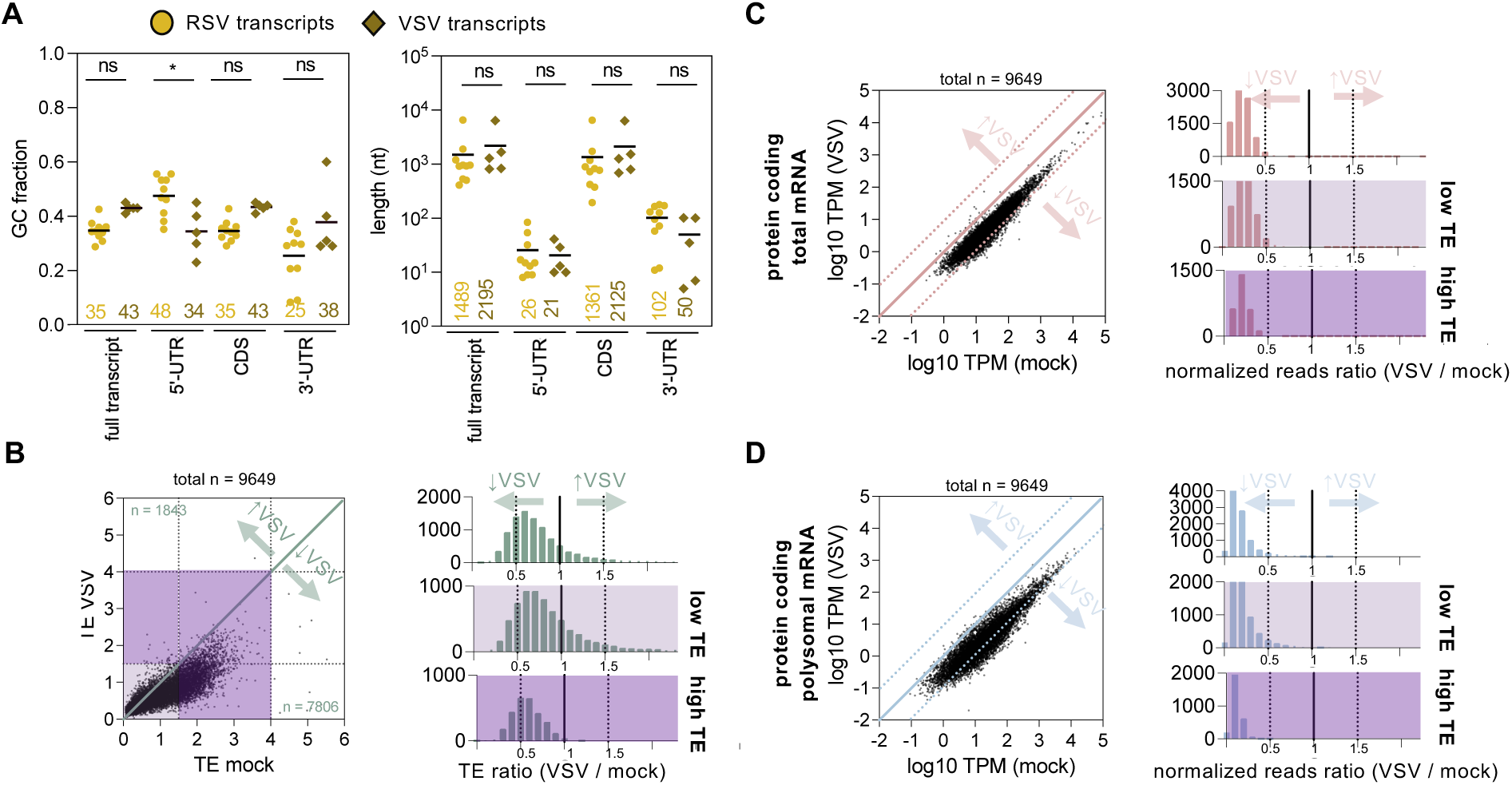
VSV infection induces the same relative enhanced ribosome recruitment for transcripts with low TE. **(A)** Distribution of GC% and length of viral protein-coding transcripts comparing RSV and VSV. Average GC% and length values are displayed underneath and shown as horizontal lines. P values were calculated with one-way ANOVA with Tukey’s multiple comparisons test (P values: * < 0.05, ns: not significant). **(B)** Translation efficiency (TE) was calculated as in Figure 3A from published dataset (Neidermyer and Whelan 2019). Scatterplots of TE comparing mock-and VSV-infected samples (MOI 10, 6h) with a global overview (*left)* and corresponding histograms *(right)* shown representing the fold-change between mock-and VSV-infected samples for all, low TE (< 1.5) and high TE (> 1.5) transcripts. **(C,D)** Scatterplots of normalized reads for total **(C)** and polysomal mRNAs **(D)** between mock-and VSV-infected samples. Histograms corresponding to the fold change between mock-and VSV-infected samples for all, low TE (< 1.5) and high TE (> 1.5) transcripts. See also **Figure S4.**

Next, to determine how the components of the TE term (*i.e.* changes in abundance or changes in polysome association) contribute to TE changes during VSV infection, we plotted normalized reads (transcripts per million – TPM) between mock-and VSV-infected samples for polysome-associated and total mRNAs. Both total and polysome-associated RNA fractions contain 60% viral reads ^47^, and therefore will include a bias due to read normalization without spike-in mRNAs. As noted above, VSV mRNAs are thought to be highly abundant and translated more than the host mRNAs. As expected, therefore, the relative abundance of host mRNAs in both total and polysomal populations was substantially decreased (**Figures 4C-D**, dots below the diagonal). Similar to RSV infected cells, changes in TE in both low and high TE subsets are caused by strong differences in polysome-associated mRNAs with no major changes in total mRNA on top of the baseline decrease due to the presence of viral mRNAs (**Figures 4C-D**, **S4B**). As described previously, to confirm our method for assaying TE, we plotted GC content ^35^ and CDS length ^36,37^ and confirm that the high TE transcripts (>1) contain significantly shorter CDSs and higher GC-content (**Figure S4C**). Overall, these findings indicate that both RSV and VSV redistribute ribosomes from high TE host mRNA towards low TE mRNAs.

### Longer AU-rich transcripts are specifically enriched in polysomes during RSV infection

To uncover transcripts that have a significantly different TE between mock-and RSV-infected samples, we performed differential expression analysis of the TE of protein-coding transcripts using DESeq2 (*i.e.* ratios of polysomal A+ mRNA to total A+ mRNA with cut-off padj < 0.05 and log2 FC ≤ -0.58 or FC ≥ 0.58, see methods) to account for any changes at the mRNA abundance level (either caused by transcription or degradation). We found several coding transcripts with significantly different TE (**Figure S5A, Table S5**, n = 533 increased and n = 46 decreased). Next, we compared features between the statistically significant cohorts and found that RSV-induced translationally upregulated transcripts have a significantly lower GC% and that translationally downregulated transcripts a significantly higher GC% compared to all coding transcripts (**Figure 5A**, full transcript, **Table S6**). To ensure that the increase of AU-rich transcripts in the polysomes is not caused by a general upregulation of AU-rich transcripts during RSV infection, we also compared the GC% of differentially abundant transcripts against all coding transcripts and found no increase of AU-rich genes (**Figure 5A**). The correlation between increased translation and lower GC% was predominantly linked to the coding sequence (CDS) and 3’-UTR and not the 5’-UTR (**Figure 5A**, **Table S6**). To confirm these RNA-seq based observations, a random cohort of highly and lowly translated transcripts were selected (**Figure S5B-C**) and validated by qRT-PCR (**Figure S5D**).

**Figure 5.**
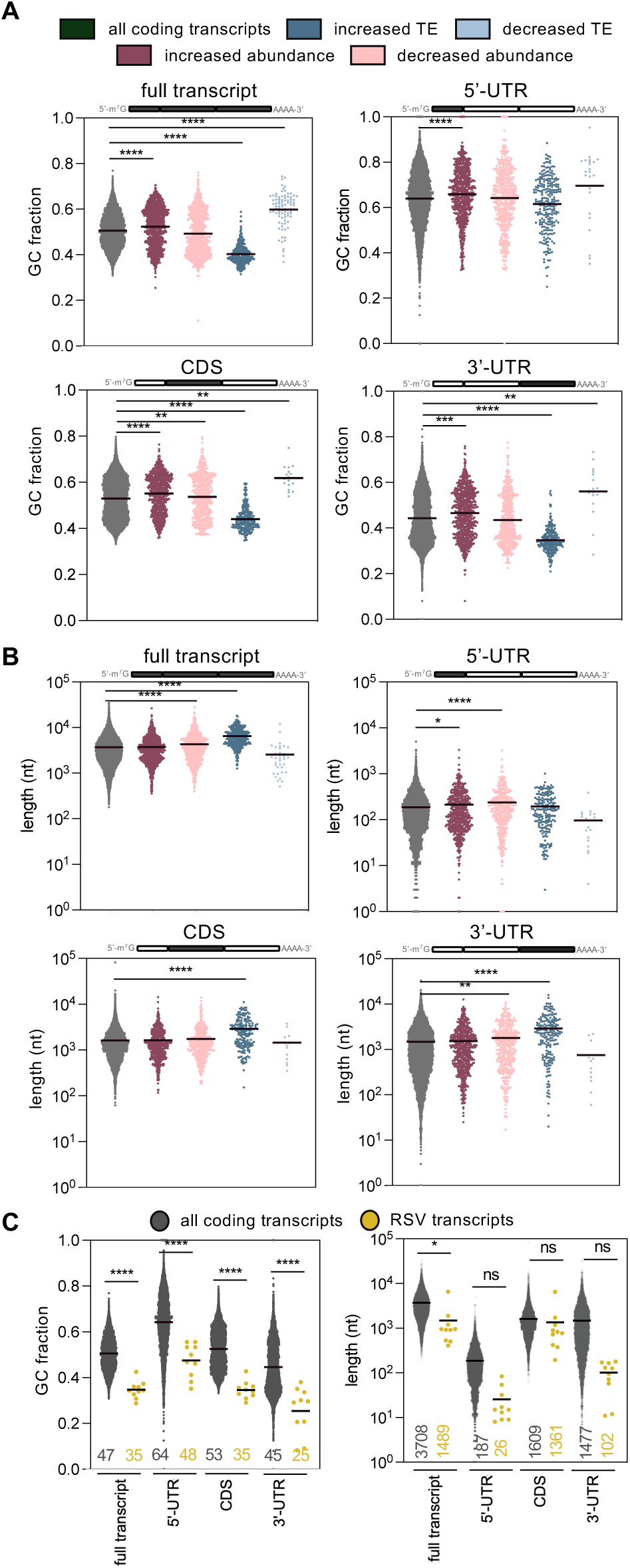
Transcripts with significantly increased TE during RSV infection are more AU-rich and contain longer CDSs and 3’-UTRs. **(A,B)** Scatterplots of GC-content (**A**) and transcript length (**B**) of host protein-coding transcripts with significantly increased or decreased abundance and TE comparing RSV-and mock-infected samples (FDR < 0.05, FC > 1.5 and FC < 1.5). P values were calculated with one-way ANOVA with Tukey’s multiple comparisons test (P values: **** < 0.0001, *** < 0.001, ** < 0.01, * < 0.05). Averages are shown as horizontal lines. **(C)** Distributions of GC-content and length of viral protein-coding transcripts compared to all host coding transcripts. Average GC% and length values are displayed underneath and shown as horizontal lines. P values were calculated with one-way ANOVA with Šídák’s multiple comparisons test (P values: **** < 0.0001, * < 0.05, ns: not significant). See also **Figure S5.**

In addition, more highly translated mRNAs during RSV infection appear to have a longer transcript length, again linked to the CDS and 3’-UTR (see **Figure 5B**, **Table S6**). To confirm that these two factors contribute independently, we confirmed that there is no correlation between GC% and transcript length (**Figure S5E**, R^2^ = 0.049). These data suggest that during RSV infection, longer AU-rich host transcripts are more efficient at recruiting ribosomes, while shorter GC-rich host transcripts are less efficient. This is consistent with the general trend we observed where transcripts with low TE, which are longer GC-poor transcripts, become more efficient at ribosome recruitment (see **Figures 3** and **4**).

### Translationally upregulated host and viral mRNAs are both AU-rich

While most host transcripts have a GC-content of 35-60%, RSV transcripts have relatively low GC-content (**Figure 5C**, GC% range from 29% to 43%), which is reflected in each of their 5’-UTR, CDS and 3’-UTR. Transcript lengths between virus and host are generally similar (**Figure 5C**, length). These observations suggest that the translational landscape of host transcripts, being biased to favor transcripts that have low GC-content during infection, may reflect an underlying trend generated during RSV infection to enhance translation of viral transcripts. A similar trend towards increased translation of lengthy AU-rich transcripts has previously been described in VSV, which like RSV, encodes 5’-capped and polyadenylated transcripts ^47^, and causes the same relative enhanced ribosome recruitment for transcripts with low translation efficiency (see **Figure 4**).

UTRs of transcripts can contain many *cis*-acting regulatory elements to either regulate translation. These include stem-loops, IRESs and upstream open reading frames (ORFs) in the 5’-UTR, as well as sequences that can be recognized by regulatory RNA-binding proteins, polyadenylation elements and the poly(A) tail in the 3’-UTR ^48^. Since RSV 5’-UTRs are very short (**Figure 5C**) and the 5’-UTR GC-content and length of differentially translated host mRNAs during RSV infection remains unchanged from the uninfected control (**Figure 5A**), the potential for regulatory elements within these sequences is relatively low. In contrast, host transcripts with 3’-UTRs that were AU-rich tended to be translated better (**Figure 5A**), similar to the case of viral transcripts (**Figure 5C**). Therefore, we focussed on elements found within the 3’-UTR of differentially translated host and viral mRNAs.

First, we compared the poly(A) tail length between differentially expressed and translated mRNAs. We used the previously published dataset which determined average poly(A) tail length ^49^, but found no statistically significant differences between translationally up-or downregulated transcripts (**Figure S5F**), which is consistent with previous findings where poly(A) tails length did not correlated with TE ^49^. Next, many RNA-binding proteins are known to specifically recognize and bind to specific conserved sequence elements ^50^ and a large number of RNA-binding proteins are known to affect translation of specific transcripts through regulatory elements found within the 3’-UTR ^51,52^. We used simple enrichment analysis (SEA) ^53^ to determine previously described RNA-binding protein motifs within the 3’-UTRs of translationally upregulated transcripts in RSV infected cells. We identified six major sequence motifs within this group of mRNAs (**Figure S5G**) and compared these RNA-binding protein motifs against motifs identified within 3’-UTRs of viral transcripts and found multiple comparable groups (**Figure S5G**). This data indicates that 3’-UTR binding host proteins could regulate a shift in AU-rich translation during viral infection.

### RSV-M2-1 binds AU-rich transcripts and associates with polysomes

While translation of AU-rich transcripts could be regulated by a host RNA-binding protein (see **Figure S5G**), viral RNA-binding proteins are also present at high levels during infection. To further investigate how AU-rich transcripts are enriched in the polysomes during infection, we investigated the role of the viral M2-1 protein which associates with all viral mRNAs ^16^. Viral mRNAs are transcribed by the viral RNA-dependent RNA polymerase within cytoplasmic inclusion bodies and M2-1 has been proposed to shuttle nascent viral mRNAs from inclusion bodies to the cytoplasm ^12^. To determine whether M2-1 is limited to bridging the mRNA to initiating ribosomes or if instead M2-1 remains associated with translating polysomes, we fractionated mock-and RSV-infected lysates by sucrose fractionation and collected fractions which were analysed by western blot. We found that M2-1 is located with both actively translating polysomes in HEp-2 and A549 cells (**Figures 6A** and **S6A-B**), the 40S ribosomal subunit and 80S ribosome (**Figure S6C**), indicating that M2-1 associates with translating polysomes during infection. We observed that another viral protein, RSV-P, is not associated with polysomes and are including this viral protein as a negative control. In addition, we observe a strong association of the nucleocapsid protein RSV-N with the heavy polysome fractions, however, RSV-N remains associated with heavy fractions following RNase treatment (see later), suggesting association with another large molecular weight complex rather than the ribosome.

**Figure 6.**
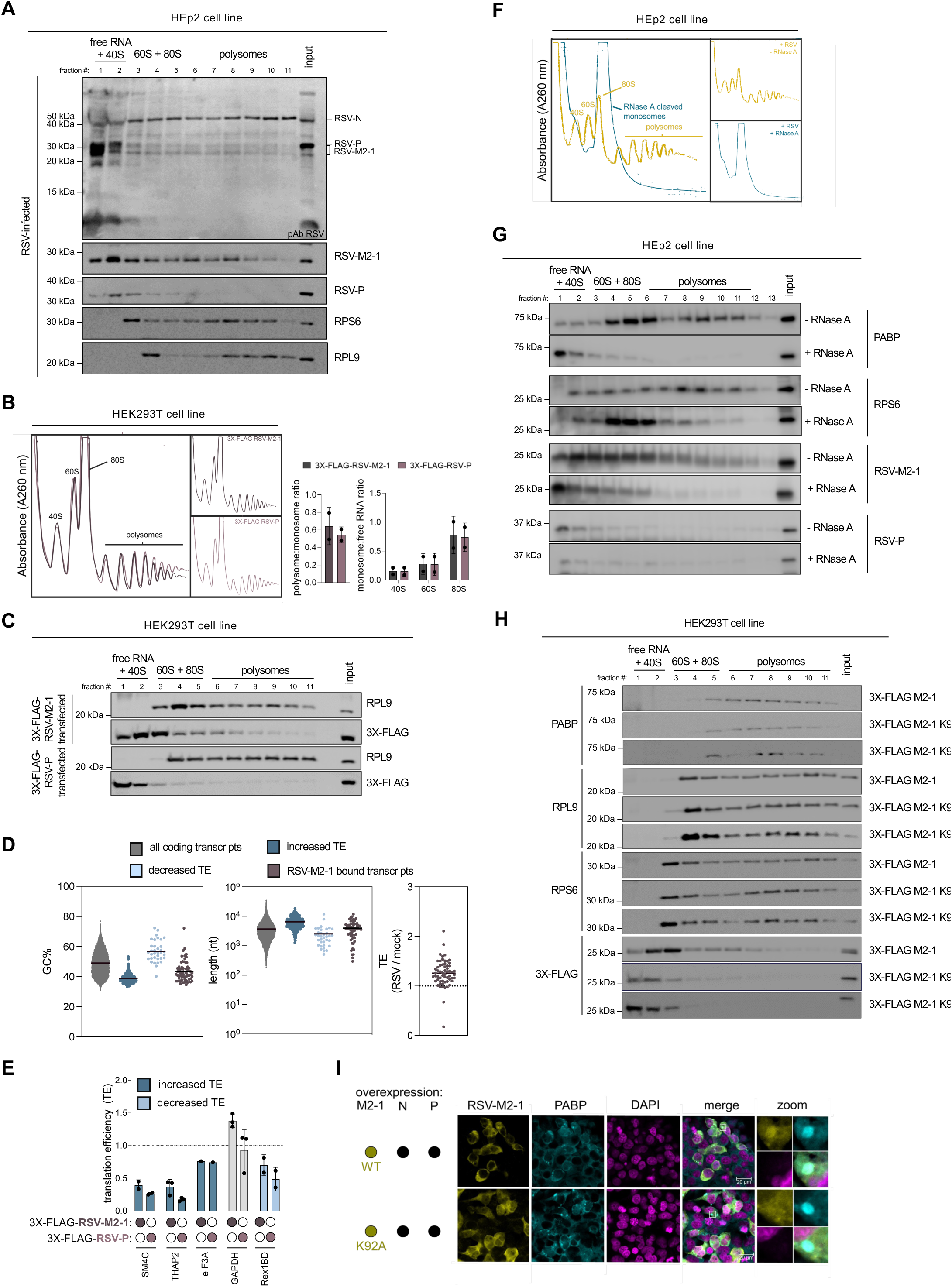
Viral M2-1 protein associates with polysomes independent of viral infection and mostly via direct mRNA-interactions. **(A)** Western blot following sucrose gradient fractionation. Fractions were collected and analyzed by western blotting for PABP, RPL9, polyclonal antibody anti-RSV and monoclonal antibodies RSV-N, RSV-P and RSV-M2-1. **(B)** Polysome profiles of HEK293T cells transfected with 3X-FLAG-M2-1 and 3X-FLAG-P (negative control) fractionated by sucrose gradient. Quantification of area under the curve calculated as in Figure 1. **(C)** Western blot of sucrose gradient fractions detecting transfected FLAG-tagged proteins from **B** using anti-FLAG antibody. RSV-M2-1 associates with polysomes without viral infection. **(D)** Distribution of GC% and transcript length of host transcripts bound by M2-1. Dataset obtained from (Braun *et al*. 2021). Averages are shown as horizontal lines. **(E)** qRT-PCR comparing translation efficiency (TE) between 3X-FLAG-RSV-M2-1 and 3X-FLAG-P (negative control) transfected HEK293T cells. TE (polysomal RNA / input RNA) for RSV / mock fold-enrichment was calculated by the ratios of ΔΔCt normalized against 5.8S rRNA. **(F)** Polysome profiles of mock-and RSV-infected HEp2 cells at MOI 1 for 24h. RNase A treatment was performed prior to loading lysates on the sucrose gradient. **(G)** Western blot following fractionation on sucrose gradients detecting RNase A-treated lysates from **F**. Fractions were collected and analyzed by blotting for direct-mRNA binding protein PABP, ribosomal core protein RPL9 and RSV proteins. **(H)** Western blot following sucrose gradient fractionation of HEK293T cells transfected with 3X-FLAG-M2-1, and poly(A) deficient binding mutants 3X-FLAG-M2-1 K92A and 3X-FLAG-M2-1 K92D. Transfected M2-1 and mutants were detected using anti-FLAG antibody. Only wild-type M2-1 associates with polysomes. **(I)** Indirect immunofluorescent staining detecting RSV-M2-1 and PABP in HEK293T cells co-transfected with inclusion body scaffolding proteins RSV-N and RSV-P. Either wild-type or mRNA-deficient M2-1 K92A mutant were co-transfected to determine co-localization. Nuclei were stained using DAPI. The white box corresponds to 10 μm and is enlarged in the zoom panel to visualize inclusion bodies. See also **Figure S6.**

Next, to determine if the viral M2-1 protein is sufficient to associate with polysomes, we transfected FLAG-tagged RSV-M2-1 and non-polysome associating RSV-P in HEK293T cells. Polysome traces between RSV-M2-1 and RSV-P transfected HEK293T cells were similar, indicating that M2-1 alone does not change overall host translation levels (**Figure 6B**). Next, polysome fractions were analysed by western blot and, consistent with viral infection, M2-1 associates with translating polysomes in HEK293T cells (**Figure 6C**). These indicates that M2-1 remains associated with the polysomes during translation even in absence of viral mRNAs, other viral factors or viral-induced host factors.

Next, we analysed a previously published RSV-M2-1 CLIP-seq dataset which described that in addition of non-specific binding to viral transcripts, M2-1 associates with a specific set of host transcripts ^16^. We determined the GC% of the M2-1-interactome and found a relatively low GC-content in comparison to all coding transcripts and more comparable to transcripts with increased TE during infection (**Figure 6D**, GC%), including specific changes in the CDS and 3’-UTR but not the 5’-UTR (**Figure S6D** and see **Figure 5A**). We found similar results for the length of these transcripts (**Figures 6D** and **S6D**, length). Comparison between the host transcripts bound by M2-1 and TE between RSV-and mock-infected samples demonstrates that most of these transcripts have a higher abundance in polysomes following RSV infection (**Figure 6D**, TE plot shows most datapoints > 1). These data indicate that M2-1 might function in recruiting ribosomes to these mRNAs.

To test if the viral M2-1 protein is sufficient to introduce a shift towards higher translation of AU-rich transcripts, we transfected 3X-FLAG-RSV-M2-1 in HEK293T cells and performed polysome profiling combined with qRT-PCR. To determine the TE of previously validated translationally up-and downregulated transcripts (see **Figure S5D**), we fractionated M2-1-and P-transfected lysates by sucrose fractionation and collected heavy polysomes. We found that the TE of the cohort of translationally upregulated transcripts yielded similar results as the translationally downregulated transcript and control GAPDH between M2-1 and P transfected cells (**Figure 6E**). This indicates that presence of M2-1, without viral infection, is not sufficient to introduce a shift towards translation of AU-rich transcripts.

### RSV-M2-1 associates with polysomes through direct mRNA interactions

Next, we determined if M2-1 is an mRNA-associated or an mRNA-independent ribosome associated protein. We treated RSV-infected lysates with RNase A prior to polysome fractionation to specifically degrade mRNA, as demonstrated previously ^54^. As a result of mRNA degradation, mRNA-associated factors (*i.e.* PABP) relocate to free RNA fractions and mRNA-independent ribosome associated proteins (*i.e.* RPS6) are found with the 80S monosomes which are minimally affected by RNase A digest (**Figure 6F**). Following RNase A treatment of RSV-infected lysates, we found M2-1 to shift from polysomes into the free RNA fractions, similarly as PABP. This suggests that M2-1 associates with polysomes through mRNA interactions (**Figure 6G**). Similarly, RNase A treatment of 3X-FLAG-RSV-M2-1 expressing HEK293T lysates (without viral infection) results in a shift of M2-1 mostly towards to free RNA fractions (**Figure S6E-G**). We performed the same polysome fractionations with transfected M2-1 K92 mutants which have lost binding affinity to mRNA ^55^ and found that M2-1 K92 mutants do not associate with polysomes anymore (**Figure 6H**). Inclusion bodies can be formed by co-transfection of RSV-P and -N (**Figure 6I**, foci in PABP staining). Interestingly, M2-1 fails to co-localize with inclusion bodies without mRNA-binding ability (**Figure 6I**, compare WT M2-1 and M2-1 K92A). These data further support the model that M2-1 associates with polysomes partially through mRNA and that co-localization with inclusion bodies might be a prerequisite for polysome association.

## Discussion

An important question for viruses that do not induce “host shutoff” is how they successfully compete with host transcripts for ribosomes required for translation of their viral transcripts. We found that RSV maintains global translation and that ribosomes are redistributed from host transcripts that are normally highly efficient at ribosome recruitment to host transcripts that are less efficient. We also found that RSV transcripts are not efficient at recruiting ribosomes and consist of AU-rich sequences. Interestingly, host transcripts with significantly increased translation efficiency (TE) were found to be longer and AU-rich, indicating that the translational landscape of host transcripts may reflect an underlying trend that is created by RSV to enhance translation of viral transcripts.

### RSV infection maintains translation and redistributes 80S monosomes into the translating pool of ribosomes

We showed that translation initiation was not inhibited through polysome profiling and instead found that polysome peaks are increased during RSV infection. This was accompanied by a decrease in 80S monosomes, indicating that monosomes are being redistributed to the polysomes. While polysome profiling is a powerful method to obtain a global overview of the distribution of 80S ribosomes compared to polysomes, it is important to note that not all transcripts found within polysomal fractions are undergoing active translation. More specifically, ribosome pausing occurs relatively frequently during translation ^56^. Rare codons, caused by low availability of matching tRNAs, are known to cause elongating ribosome to pause ^57,58^. Ribosomal pausing could be utilized by RSV to promote co-translational folding of viral proteins or to enhance endoplasmic reticulum (ER)-targeting of viral membrane proteins RSV-G, -F and -SH. Additionally, viruses often employ ribosomal pausing through a slippery sequence to induce programmed ribosomal frameshifts to enhance their coding capacity ^59^. However, when ribosomes undergo prolonged pausing, the potential for ribosomes collisions exists which leads to formation of ribosomes complexes containing two (disomes), three (trisomes) or more ribosomes ^60^. While ribosome collisions are eventually resolved by several surveillance pathways ^60^, this could result in larger polysomes. More specifically, treatment with intermediate concentrations of translation elongation inhibitors (such as anisomycin), leads to increased ribosome collision and has been found to decrease 80S monosomes and increase polysomes ^61^, similar to our polysome traces comparing mock-and RSV-infected cells.

The number of ribosomes found within polysomes is determined by a combination of translation initiation and elongation rates, where faster translation initiation and slower translation elongation both enhance polysome formation ^62,63^. In addition, larger polysome peaks can also be induced by increased polysome-association by transcripts with longer CDSs which can accommodate more ribosomes. It has indeed been shown that the number of ribosomes associated with a transcript correlates with the CDS length ^64^ and that transcripts with short ORFs (< 500 nt) are typically found more frequently as 80S monosomes as opposed to polysomes ^62^. We found that during RSV infection transcripts with longer CDSs are indeed specifically enriched in polysomes which could contribute to the observed increased polysome peaks.

### Low ribosome occupancy transcripts are longer and AU-rich and become more efficient at ribosome recruitment during infection

While previous work has shown that viral proteins alter host transcription through direct chromatin interactions ^65,66^, limited information on translational changes have been described to date. We found that during RSV infection, ribosomes get redistributed from transcripts that are normally efficient at ribosome recruitment to transcripts that are less efficient. This redistribution could benefit the virus for several reasons. First, we found that viral mRNAs are highly abundant in the total RNA fraction (14%), but are present in low numbers in the polysomes (2%) indicating that RSV mRNAs have relatively low TEs. Viral mRNAs are produced within cytoplasmic inclusion bodies where they accumulate before being released into the cytoplasm for translation ^10–12^ which could partially contribute to the observed low TE of viral transcripts. With less viral transcripts accessible to ribosomes, the virus could benefit from a global shift in higher ribosome recruitment for low TE transcripts. Second, RSV transcripts contain very short 5’-UTRs. More specifically 7 out of 10 viral transcripts contain a 5’-UTR shorter than 20 nucleotides, which has been linked to less efficient ribosome recruitment ^67^. Besides a decreased efficiency in ribosome recruitment, another consequence of short 5’-UTRs is that translation initiation can occurs at a downstream start codon as opposed to the 5’-cap proximal start codon ^68^. This could result in non-canonical protein production of shorter viral proteins or completely novel proteins in a different frame which could affect immune response pathways ^69^.

A possible cause for the global redistribution of ribosomes from high to low TE transcripts is mature tRNA level availability. More specifically, decreased availability of specific mature tRNAs could decrease translation for a selection of transcripts ^70^ since translation elongation slowdowns decrease translation initiation rates ^71–74^. A recent study however has found that while mature tRNA levels are different during differentiation, the tRNA anticodon pool remains the same which maintains the decoding speed of elongating ribosomes ^75^. Similarly, it has been shown that while large differences exist in isodecoder expression in different tissues, the anticodon pool remains similar ^76^. It remains to be determined if these rules are also valid during viral infections and if RSV induces changes in mature tRNA levels that could cause these global ribosome redistributions.

### M2-1 associates with polysomes and AU-rich transcripts with increased translation efficiency

We found that transcripts with a significantly increased TE contain shorter and more AU-rich 3’-UTR sequences. Since many RNA-binding proteins can affect translation of specific transcripts through regulatory elements found within the 3’-UTR ^51,52^, we identified common sequence motifs between viral and translationally upregulated transcripts. Activation or upregulation of one of these translationally enhancing RNA-binding proteins could result in specific enhanced translation enrichment for transcripts containing the corresponding 3’-UTR binding site.

In addition to host proteins, viral RNA-binding proteins are also highly abundant during infection. We found that during infection, RSV-M2-1 associates with both the 40S subunit, 80S monosomes and translating polysomes. This interaction was found to be mainly through mRNA-interactions. Consistent with this is previous work demonstrating that M2-1 directly interacts with PABP ^77^. This could indicate that M2-1 could function as an important component to enhance translation initiation of the bound mRNA. More recently, viral polysome associated proteins have been identified in other viruses. For example, VP22 was identified in herpes simplex virus-1 (HSV-1) where it associated with both initiating and elongating ribosomes ^78^.

Since M2-1 can associate with polysomes in HEK293T cells in absence of viral infection, we hypothesized that M2-1 could function in recruiting ribosomes of not only viral transcripts, but also AU-rich host transcripts. We tested this by transfection of M2-1 in HEK293T cells and determined if this overexpression of M2-1 would be sufficient to induce a shift towards translation of AU-rich transcripts. While we could not validate this hypothesis, it is possible that M2-1 plays an essential role in this process even if it is not sufficient to drive AU-rich translation in isolation. An important difference between RSV-infected cells and M2-1 transfected cells is the absence of inclusion bodies in the latter. These are important membraneless subcellular compartments in which viral replication and transcription take place. Previous work demonstrated that M2-1 binds newly transcribed viral transcripts within these inclusions bodies and shuttles these into the cytoplasm for translation ^12^. Another argument for the importance of inclusion bodies in regulating the AU-rich translational shift is that during vesicular stomatitis virus (VSV) infection, host reporter genes containing flanking gene-start and gene-end viral sequences have enhanced translation, however only when these sequences are inserted into the viral genome as opposed to expression from a DNA plasmid ^79^. This indicates that transcription is an important determinant of translation efficiency in VSV infected cells. In addition to inclusion bodies, many other changes occur in RSV infected cells including many transcripts that have increased translation, resulting in higher expression of their corresponding protein which could also have a role in assisting in specific recruitment of AU-rich transcripts to polysomes.

## Acknowledgements

We thank Dr. Ultan Power (Queen’s University Belfast, UK) for providing us with HEp-2 and A549 cell lines used in this study. We thank members of the Bayfield and Guydosh lab for feedback on this project. This research was supported by a Project Scheme Grant from the Canadian Institutes of Health Research’s Institute of Genetics (application #419907 to M.A.B.) and the Intramural Research Program of the NIH, The National Institute of Diabetes and Digestive and Kidney Diseases (NIDDK) (DK075132 to N.R.G.).

## Author contributions

Conceptualization, K.K., N.R.G. and M.A.B.; Methodology, K.K.; Investigation, K.K.; Writing – Original Draft, K.K.; Writing – Review & Editing, N.R.G. and M.A.B.; Funding Acquisition, N.R.G. and M.A.B; Resources, N.R.G. and M.A.B.; Supervision, N.R.G. and M.A.B.

## Data availability

High-throughput RNA sequencing has been deposited to the Gene Expression Omnibus (GEO) under the accession number GSE268742.

## Supplemental information

Document S1. Figures S1-S6 and Table S8.

Document S2. Tables S1-S7. Excel file containing additional data too large to fit in a PDF.

## Materials and Methods

### Cell culture, RSV infection and arsenite treatment

HEp-2 cells were grown in DMEM containing 5% FBS. HEK293T and A549 cells were grown in DMEM containing 10% FBS. Cell lines were maintained in a humidified incubator at 37°C with 5% CO_2_. Arsenite (NaAsO_2_) treatment was done by incubating cells with 0.5 mM (unless otherwise stated) for 1 hour at 37°C.

The RSV strain A2 (ATCC, serial passage-1) was propagated in HEp-2 cells. Briefly, 15-cm plates at 80% confluency were infected with RSV (P2, MOI = 0.1) for 2 hours at 37°C in 5 mL FBS-free DMEM. Following infection, the cells were maintained in DMEM containing 1% FBS and incubated for approximately 3 days until syncytia formed. The cells were scraped, and supernatant was collected following centrifugation at 1000g for 15 minutes at 4°C. The RSV stock P3 was aliquoted, snap frozen in liquid nitrogen and stored at -80°C. Titration of the RSV stock was performed according to the Tissue Culture Infectious Dose-50 (TCID_50_) Spearman–Kärber method ^80^.

For experiments, RSV infections were done using the titrated RSV P3 stock. In brief, cells were grown overnight and the RSV P3 stock, quickly thawed at 37°C and diluted in FBS-free DMEM to the desired MOI. The cells were washed once with PBS, followed by incubation with a small volume of FBS-free DMEM (*i.e.* 15 cm plates: 5 mL, 10 cm plates: 2 mL, 24-well plates: 200 μL, 96-well plate: 32 μL) and incubated for 2 hours with frequent rocking to redistribute the infection medium evenly. Mock treatment included, PBS wash and 2 hour incubation in infection medium. Following infection, the cells were maintained in DMEM containing 5% (HEp-2) or 10% inactivated FBS (A549) (30 minutes at 56°C) for 24 hour unless stated otherwise.

### Polysome profiling

Polysome profiling was performed as described in ^81^. In brief, 100 μg/mL cycloheximide was added to the cells for 5 minutes at 37°C prior to collection. The cells were washed twice in PBS containing 100 μg/mL cycloheximide. The cell pellets were stored at -80°C until use. Next, the cell pellets were lysed in 485 μL hypotonic buffer [5 mM Tris-HCl (pH 7.5), 2.5 mM MgCl2, 1.5 mM KCl, 1X Halt™ Protease Inhibitor Cocktail, 100 μg/mL cycloheximide, 2 mM DTT, 200 Units/mL SUPERase In™ RNase Inhibitor, 0.5% (v/v) Triton X-100 and 0.5% (w/v) sodium deoxycholate], followed by centrifugation for 5 minutes at 20,000g at 4°C to obtain cytoplasmic extracts. A fraction of the lysate was taken as the total protein samples. The remaining sample (500 μL of 20 A260 units) was fractionated on a 7-step 20-50% sucrose gradient prepared in sucrose buffer [20 mM HEPES (pH 7.6), 100 mM KCl, 5 mM MgCl2 and 100 μg/mL cycloheximide] by centrifugation at 30,000 RPM for 3 hours in a Beckman SW41Ti rotor at 4°C (acceleration: max., deceleration: no brake). Polysome profiles were obtained with BRANDEL Density Gradient Fractionation System by measuring the absorbance at 254 nm with the UA-6 Detector in a continuous flow. Polysome fractions were collected (800 μL each, fraction numbers 1-11), unless otherwise stated. Polysome traces were obtained with the build-in Chart Recorder with paper and pen and digitally represented using Inkscape v.1.2.1.

RNase A treated samples are processed similarly as above, with a few changes. Cell pellet lysis occurs in hypotonic buffer omitting RNase inhibitor [5 mM Tris-HCl (pH 7.5), 2.5 mM MgCl2, 1.5 mM KCl, 1X Halt™ Protease Inhibitor Cocktail, 100 μg/mL cycloheximide, 2 mM DTT, 0.5% (v/v) Triton X-100 and 0.5% (w/v) sodium deoxycholate]. Cell lysates (500 μL of 20 A260 units) were treated with 6 ng/uL RNase A for 30 minutes at room temperature, followed by addition of 200U SUPERase In™ RNase Inhibitor.

RNA extraction of polysome fractions was performed by adding 2 parts 100% ethanol containing 80 mM NaOAc, pH 5.1 and 300 μg GlycoBlue overnight at -80°C to precipitate the RNA. RNA pellets were collected by centrifugation at 20,000g for 30 minutes at 4°C, followed a 70% ethanol wash and resuspension in ddH_2_O. Both polysomal RNA and total RNA were extracted using Trizol according to manufacturer’s instruction.

Protein extraction of polysome fractions was done by incubation in 10% Trichloroacetic acid (TCA) overnight at -20°C to enhance protein precipitation, followed by centrifugation for 15 minutes at 10,000g at 4°C. The protein pellet was washed twice with ice-cold 100% acetone, air-dried overnight and resuspended in 150 μL/mL sucrose gradient 2.5X Laemmli buffer [5X SDS loading dye: 5% β-mercaptoethanol (v/v), 0.02% bromophenol blue (w/v), 30% glycerol (v/v), 10% sodium dodecyl sulfate (SDS) (w/v), 250 mM Tris-HCl, pH 6.8].

### Western blot

Cellular lysates were quantified using the Pierce Coomassie Plus (Bradford) Assay Reagent to obtain protein concentrations. Protein samples were incubated with 1x Laemmli buffer [5X Laemmli buffer: 5% β-mercaptoethanol (v/v), 0.02% bromophenol blue (w/v), 30% glycerol (v/v), 10% sodium dodecyl sulfate (SDS) (w/v), 250 mM Tris-HCl, pH 6.8] for 10 minutes at 95 °C and separated using a 12% SDS-PAGE for 1 hour at 110 V. Proteins were transferred to a nitrocellulose membrane for 2 hours at 50 V. The nitrocellulose membrane was blocked in 5% nonfat dried milk (NFDM) in tris-buffered saline containing 0.1% Tween-20 (TBS-T) for 1 hour at room temperature or overnight at 4°C. The membrane was probed with appropriate primary antibodies in TBS-T for 1 hour at room temperature or overnight at 4°C. After primary antibody binding, the membrane was washed 5 times in TBS-T, incubated with appropriate HRP-coupled secondary antibody for 1 hour at room temperature and washed 5 times in TBS-T. Membranes were incubated with Pierce® ECL Western Blotting Substrate and imaged on a MicroChemi chemiluminescence system (DNR Bio-Imaging Systems). Secondary antibodies used were horse anti-mouse IgG HRP (1:10,000), goat anti-rabbit IgG HRP (1:10,000) and Rabbit Anti-Goat IgG HRP (1:10,000). Western blot quantifications were done using Image J v2.14.0.

In order to probe the same membrane for proteins with similar molecular weight with multiple primary antibodies raised in different species (*e.g.* rabbit and mouse) we performed mild stripping of the western blot membranes to quench HRP. In brief, the membrane was incubated twice in stripping buffer-HCl, pH 2.2 [1.5% glycine (w/v), 0.1% SDS, 1% Tween-20] for 10 minutes, followed by two 5-minute washes with PBS and two 5 minute washed with TBS-T. Prior to probing with primary antibody, the membrane was blocked as described above.

### RT-qPCR

Total RNA was extracted using Trizol extraction (according to manufacturer’s instructions). The isolated RNA was resuspended in 1X Reaction Buffer with 2 units of Turbo DNase and incubated for 30 minutes at 37°C. The DNase-treated samples were subsequently Trizol again extracted to inactivate DNase activity. Next, 50 ng/μL DNase-treated RNA was reverse transcribed using the iScript cDNA Synthesis Kit according to manufacturer′s instructions. The cDNA was diluted to 25 ng (in 12.5 μL per technical triplicate) and quantified using the SensiFAST SYBR No-Rox kit with 12.5 μM of each primer (1x forward, 1x reverse) (**Table S8**) using following qPCR settings: 95°C for 5 minutes and 35 cycles of 5 sec at 95°C and 15 sec at 60°C, followed by a melting curve analysis up to 99°C to confirm amplification of a single amplicon. Fold enrichment was calculated using the ΔCt method (Ct _5.8S rRNA_ – Ct _RNA_). Translation efficiency (polysomal RNA / input RNA) RSV / mock fold enrichment was calculated by the ratios of ΔΔCt normalized against 5.8S rRNA.

### DNA plasmid transfections

Viral genes were amplified from Geneblocks (IDT) (**Table S7**) using forward primers containing a 3X-Flag sequence and reverse primers contained a stop codon (**Table S8**). Next, the amplified insert was cloned into the pEGFP-N1 plasmid using SalI and BamHI restriction sites resulting in a CMV-3X-Flag-viral-gene-stop construct. Transfection of HEK293T cells was done using PolyJet according to the manufacturer’s protocol for 48h.

### Indirect immunofluorescent staining

The intracellular localization of endogenous and exogenously expressed proteins was determined through indirect immunofluorescent staining. Prior to fixing, monolayers were washed twice with PBS. Next, cells were fixed for 20 minutes with 4% paraformaldehyde in PBS and permeabilized for 10 minutes with 0.1% Triton X-100 in PBS. Next, the cells were blocked for 1 hour in 1% Bovine Serum Albumin (BSA) in PBS, followed by incubation with primary antibodies in blocking solution for 1 hour at room temperature or overnight at 4°C. Cells were subsequently washed 4 times with PBS, incubated with appropriate fluorochrome-bound secondary antibodies for 1 hour at room temperature and washed 4 times with PBS. Cells were stained for 2 minutes with 2.5 μg/mL 4’,6-diamidino-2-phenylindole (DAPI), washed twice with PBS and overlaid with 1,4-diazabicyclo[2.2.2]octane (DABCO). All images were acquired by the LSM700 laser scanning confocal microscope (Zeiss) with a 63x oil immersion objective or 20x objective. Primary antibodies used were rabbit anti-PABP (Abcam ab21060), mouse anti-G3BP (BD Biosciences, 611126), goat anti-RSV (Virostat, 601), and mouse anti-M2-1 (Abcam ab94805). Secondary antibodies used were donkey anti-rabbit IgG Alexa Fluor 594 (1:1000), goat anti-rabbit IgG Alexa Fluor 488 (1:1000), donkey anti-mouse IgG Alexa Fluor 546 (1:1000), and donkey anti-goat IgG Alexa Fluor 488 (1:1000). Immunofluorescent images were pseudo-colored: (1) blue-emitting fluorescent DAPI to magenta, (2) RSV-specific staining (red or green-emitting) to yellow and (3) host proteins (red or green-emitting) to cyan.

### Next generation sequencing and sample quality control

Following Trizol extraction of total and polysomal RNA (see polysome profiling methods section), 5 μg RNA was heated in 1X formamide [2X formamide: 95% deionized formamide, 0.025% (w/v) bromophenol blue, 0.025% xylene cyanol (w/v), 5 mM EDTA, pH 8.0] for 3 minutes at 95°C, and immediately snap cooled on ice for 3 minutes. Next, the RNA was separated on a 1% agarose gel in 1X tris-borate-EDTA (TBE) buffer for 40 minutes at 100V. Bioanalyzer data quality control of RNA samples performed at TCAG, Hospital for Sick Children, showed an RNA integrity number (RIN) of 10 for all samples.

RNA extracted from total and polysomal RNA (see polysome profiling methods section) was subjected to stranded cDNA library preparation by poly(A) tail selection (poly(A)+) (NEBNext) and paired end 50 bp sequencing using the Illumina NovaSeq 6000 at The Centre for Applied Genomics (TCAG, Hospital for Sick Children). Raw reads in .fastq format were trimmed using Trim Galore v.0.5.0 with Cutadapt v.1.10 (with following parameters: -q 25, --clip_R1 6, --clip_R2 6, --stringency 5, --length 40, --paired). Quality control was done using FastQC v.0.11.5 before and after trimming. The raw trimmed reads were aligned to the concatenated GRCh38 GENCODE release 36, and RSV genome (GenBank: KT992094.1) using STAR aligner v.2.6.0c ^82^. Gene expression analysis was done using htseq-count v.0.6.1p2 (mode “intersection_nonempty”) to obtain raw counts (**Table S1**). Raw counts were used for differential expression analysis using DESeq2 v.1.32.0 to obtain DESeq2-normalized counts through the median of ratios method (*i.e.* normalized for sequencing depth and RNA composition) (**Tables S2-S3**) ^33^. The DESeq2 design matrix contained information for the component *virus* (distinguishing between mock-and RSV-infection) and *RNA* (distinguishing between poly(A)+ total RNA and poly(A)+ polysomal RNA). Differential expression analysis comparing total and polysomal RNA between mock-and RSV-infected used the design formula *design = ∼virus* (**Tables S2-S3**). While differential expression analysis comparing translation efficiency (TE; polysomal read counts / total read counts) between mock-and RSV-infected used the design formula *design = ∼RNA + virus + RNA:virus* which takes the TE ratio into consideration (**Table S5**). Reproducibility between biological replicates was determined through multidimensional scaling (MDS) using R package limma v.3.48.0 (**Figure S2C**) and through calculating the Euclidean distance of the gene expression matrix from different samples plotted on a heatmap using R package pheatmap v.1.0.12 (**Figure S2D**).

Scatterplots displaying normalized reads and volcano plots were generated using R package ggplot2 v.3.4.1. Pie charts, one-dimensional scatterplots displaying normalized reads and two-dimensional scatterplots displaying TE, histograms, cumulative histograms, one-dimensional scatterplots displaying transcript’s GC% and length (nt), and bar graphs were generated in GraphPad Prism v.10.2.2.

### 5’-UTR, CDS and 3’-UTR analysis

Transcript sequences, including cDNA sequences (*i.e.* full length), 5’-UTR, CDS and 3’-UTR, were downloaded from Ensembl (Ensembl Genes 111; Human genes GRCh38.p14; https://useast.ensembl.org/biomart/martview). Transcripts were selected from the Matched Annotation from the NCBI and EMBL-EBI (MANE) to obtain information for representative transcripts within the human transcriptome ^83^. The RSV genome was downloaded from NCBI (GenBank: KT992094.1, https://www.ncbi.nlm.nih.gov/nuccore/KT992094.1) and the VSV genome from NCBI (GenBank: OR921183_1, https://www.ncbi.nlm.nih.gov/nuccore/2635771998). GC% and length were calculated for the MANE, RSV and VSV transcripts using a custom Python script (**Table S6**).

Simple enrichment analysis (SEA) ^53^ against the RNA motif database (Ray2013 Homo sapiens) was used to uncover RNA-binding protein motifs within the 3’-UTRs of MANE and RSV sequences.

## Supplemental Information

**S1 Related to Figure 1. RSV infection does not induce stress granule formation in HEp2 and A549 cells.**

**(A,B)** Western blot demonstrating lack of eIF2α phosphorylation during RSV infection by comparing eIF2α-P and total eIF2α levels between (**A**) mock-and RSV-infected (MOI 1, 24h) and (**B**) untreated and NaAsO_2_-treated (positive control) (0.5 mM, 1h) A549 cells. Relative quantification against control cells is shown below. RSV infection was confirmed by immunoblotting with a polyclonal anti-RSV antibody (pAb).

**(C)** Western blot comparing eIF2α-P and total eIF2α levels between mock-and RSV-infected (MOI 1) at different time points. Viral proteins were detected using a polyclonal anti-RSV antibody.

**(D,E)** RSV infection does not induce stress granule formation seen by indirect immunofluorescent staining of mock-and RSV-infected cells (MOI 1, 24h) detecting stress granule markers PABP (**D**) and G3BP (**E**). RSV proteins were detected using a polyclonal anti-RSV antibody (shown in yellow) and nuclei were stained using DAPI (magenta). The white box corresponds to 10 μm and is enlarged in the zoom panel to visualize inclusion bodies where nascent viral transcripts are transcribed.

**(F)** Indirect immunofluorescent staining of arsenite-treated cells (positive control) (0.5 mM, 1h) detecting stress granule markers PABP and G3BP. Nuclei were stained using DAPI (magenta)

**(G)** Polysome profiles of sucrose gradient fractionated mock-and RSV-infected A549 cells (MOI 1, 24h). Quantification of area under the curve between polysomes and monosomes (40S, 60S and 80S) are plotted to estimate translation levels. Quantification of area under the curve between free RNA fraction (not shown) and 40S, 60S and 80S are plotted to determine changes in free monosomes and 80S subunits.

**S2 Related to Figure 2. Quality control RNA-seq samples.**

**(A)** Western blot of total cytoplasmic protein obtained from samples used for high-throughput sequencing immunoblotted with polyclonal antibody (pAb) anti-RSV and monoclonal antibodies anti-RSV-N, anti-RSV-P and anti-RSV-M2-1 to confirm viral infection and loading control GAPDH.

**(B)** Agarose gel to determine RNA quality of RNAseq samples. Note the absence of tRNAs in the polysomal RNA.

**(C)** Multidimensional scaling (MDS) to determine similarity between RNAseq replicates. Diversity between samples are delineated by RNA type (dimension 1; total vs polysomal A+ RNA) and infection status (dimension 2; -RSV vs +RSV).

**(D)** Heatmap demonstrating reproducibility between biological replicates. Color gradient shown on the heatmap corresponds to the Euclidian distance which was calculated for gene expression matrixes and compared between samples. Biological replicates are similar in distance and cluster together.

**S3 Related to Figure 3. TE quality control data for RSV data.**

**(A)** Scatterplots TE comparing mock-and RSV-infected samples (MOI 1, 24h) for biological triplicates with a global overview (*left)* and zoomed versions (*middle and right*).

**(B)** Distribution of the GC% and CDS length of host protein-coding transcripts divided between high TE (>2) and low TE (<2). P values were calculated with an unpaired t test (P values: **** < 0.0001). Highly translated mRNAs are shorter and GC-rich.

**S4 Related to Figure 4. TE quality control data for VSV data.**

**(A)** Cumulative histograms of TE ratios (polysomal A+ RNA / total A+ RNA) to determine high vs. low TE cut-offs. Data to determine TE cut-off was derived from uninfected samples for both the RSV dataset (this paper, displayed in **Figure 3**) and VSV dataset (Neidermyer and Whelan 2019, displayed in **Figure 4**). The RSV dataset was divided between high and low TE transcripts by setting the cut-off value at 2. This approximately separates the top 32% most highly translating transcripts (TE>2) from the other 68% transcripts with low TE (>2). On the other hand, setting a TE cut-off value of 2 for the VSV dataset would result in a division of 89% (high TE) vs. 12% (low TE). Since this would likely result in a non representative dataset, we set the TE cut-off for the VSV dataset at 1 which divides the reads 50-50.

**(B)** Scatterplots of normalized reads for total cytoplasmic mRNAs (*top*) and polysome-associated mRNAs (*bottom*) between mock-and VSV-infected samples (MOI 10, 6h) from published dataset by Neidermyer and Whelan 2019. Corresponding histograms are shown in **Figures 4C,D** with light purple (low TE) and dark purple(high TE) backgrounds.

**(C)** Distribution of the GC% and CDS length of host protein-coding transcripts divided between high TE (>1) and low TE (<1) from previously published dataset from Neidermyer *et al*. 2019. P values were calculated with an unpaired t test (P values: **** < 0.0001). Highly translated mRNAs are shorter and GC-rich.

**S5 Related to Figure 5. Total and polysome-associated differentially expressed protein-coding transcripts.**

**(A)** Volcano plots of differentially expressed protein-coding host mRNAs comparing mock-and RSV-infected samples (MOI 1, 24h) (three biological replicates) for translation efficiency (polysomal vs total A+ mRNA) *(left)* and total A+ mRNA (*right,* same as **Figure 2D**, *left*). Differential abundance was calculated as the ratio of total A+ RNA in RSV-and mock infected cells and differential TE as the ratios of polysomal to total RNA between RSV-and mock-infected cells. The horizontal line indicates a cutoff of padj < 0.05 and vertical lines indicate a 1.5-fold change (FC).

**(B)** GC% and transcript length from the random cohort of highly and lowly translated transcripts transcripts confirmed in **C** and **D**.

**(C,D)** A selection of transcripts from the RNAseq dataset in (**C**) validated by qRT-PCR in (**D**). Translation efficiency (polysomal RNA / input RNA) for RSV / mock fold enrichment was calculated by the ratios of ΔΔCt normalized against 5.8S rRNA.

**(E)** Scatterplots demonstrating no correlation between GC% and transcript length.

**(F)** Distribution of the poly(A) tail length of host protein-coding transcripts with significant (FDR < 0.05) increased or decreased abundance (FC > 1.5 and FC < 1.5) comparing RSV-and mock-infected samples. P values were calculated with one-way ANOVA with Tukey’s multiple comparisons test (P values: **** < 0.0001). Dataset obtained from previously published study by Chang *et al*. 2014.

**(G)** Simple enrichment analysis (SEA) of motifs found within the 3’-UTR of statistically significantly translationally upregulated (FDR < 0.05, log2 FC > 0.58) protein-coding transcripts (*left*) compared to the 3’-UTR of viral transcripts (*right*).

**S6 Related to Figure 6. M2-1 associates with the 40S subunit, 80S monosome and polysomes independent of infection**

**(A)** Western blot of the mock-infected control from **Figure 6A**.

**(B)** Western blot confirmation in another cell line. Same experiment as in **Figure 6A**.

**(C)** Same experiment as in **Figure 6A** with higher resolution around 40S, 60S, 80S and light polysomes by collecting more fractions for 40S, 60S, 80S and light polysome fractions.

**(D)** Distribution of the GC% (*top*) and transcript length (*bottom*) of 5’-UTR, CDS and 3’-UTR of host protein-coding transcripts with significant (FDR < 0.05) increased or decreased translation efficiency (TE) during RSV infection (padj < 0.05, FC > 1.5 and FC < 1.5). FC: fold change.

**(E)** Polysome profiles of HEK293T cells transfected with 3X-FLAG-M2-1 and 3X-FLAG-P (non-polysome associating negative control). RNase A treatment was performed prior to loading lysates on the sucrose gradient.

**(F)** Western blot following sucrose gradient fractionation detecting transfected FLAG-tagged proteins from **S6E** using anti-FLAG antibody. Fractions were collected and analyzed by western blotting for direct-mRNA binding protein PABP, ribosomal core protein RPL9 and polyclonal anti-RSV antibody.

**(G)** Western blot comparing input samples from **S5F** (*top*) and **6H**.

**Table S1. RNA-seq raw counts from this study, Related to Figure 2**. Columns A-E contain gene information, columns F-K mock-infected and columns L-Q RSV-infected (MOI 1, 24h) raw counts. Total indicates total A+ RNA and pol indicates polysome-associated A+ RNA. A+: poly(A)-tail enriched.

**Table S2. DESeq2 normalized counts and differential expression for total A+ RNA from this study, Related to Figure 2**. Columns A-D contain gene information, columns E-J DESeq2 outputs for RSV / mock comparisons for total A+ RNA and columns K-P normalized total A+ RNA counts. A+: poly(A)-tail enriched.

**Table S3. DESeq2 normalized counts and differential expression for polysome-associated A+ RNA from this study, Related to Figure 2**. Columns A-D contain gene information, columns E-J DESeq2 outputs for RSV / mock comparisons for polysome-associated A+ RNA and columns K-P normalized polysome-associated A+ RNA counts. A+: poly(A)-tail enriched.

**Table S4. Translation efficiency (TE) data for all transcripts from this study, Related to Figure 3**. DESeq2 normalized counts were used to calculate translation efficiency ratios for mock-and RSV-infected samples (polysomal A+ mRNA / total A+ mRNA). A+: poly(A)-tail enriched.

**Table S5. DESeq2 normalized counts and differential expression for translation efficiencies (TEs) from this study, Related to Figure 5**. Columns A-D contain gene information, columns E-J DESeq2 outputs for translation efficiency (TE) ratios for RSV-vs. mock-infected samples (TE: total A+ RNA / polysome-associated A+ RNA) and columns K-T normalized total and polysome-associated A+ RNA counts (same as in Tables S2-S3).

**Table S6. GC% and length data for MANE selected transcripts from this study, Related to Figures 4 and 5**. GC% and length for cDNA sequences (*i.e.* full length), 5’-UTR, CDS and 3’-UTR. Transcripts were selected from the Matched Annotation from the NCBI and EMBL-EBI (MANE) to obtain information for representative transcripts within the human transcriptome.

**Table S7. Gene blocks and plasmids used in this study. Related to Methods.** 3X-FLAG sequences are underlined.

**Table S8.**
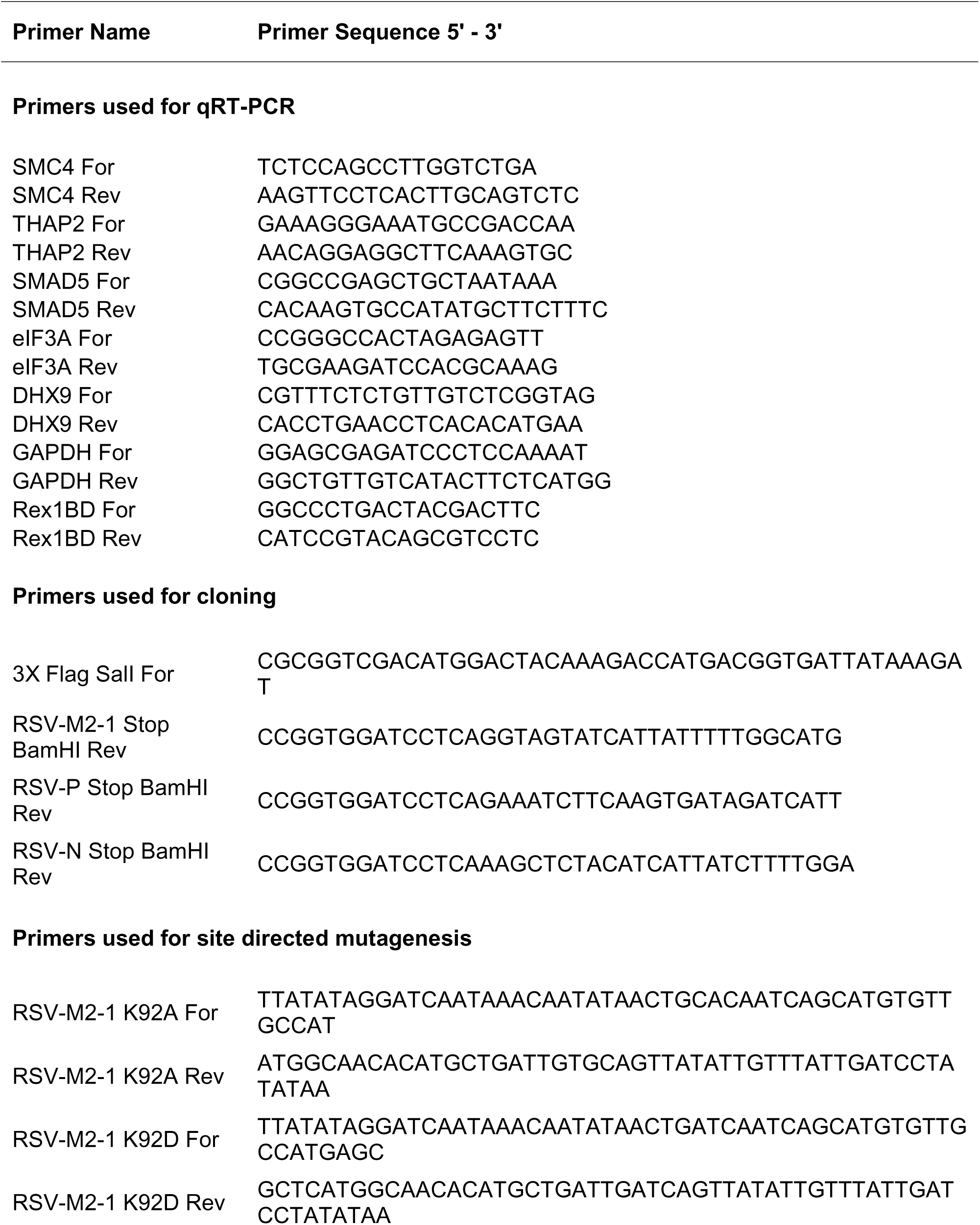
Oligonucleotides used in this study. Related to Methods.

