## Supplementary figures and images for "Respiratory Syncytial Virus (RSV) optimizes the translational landscape during infection"

### Supplementary Figures

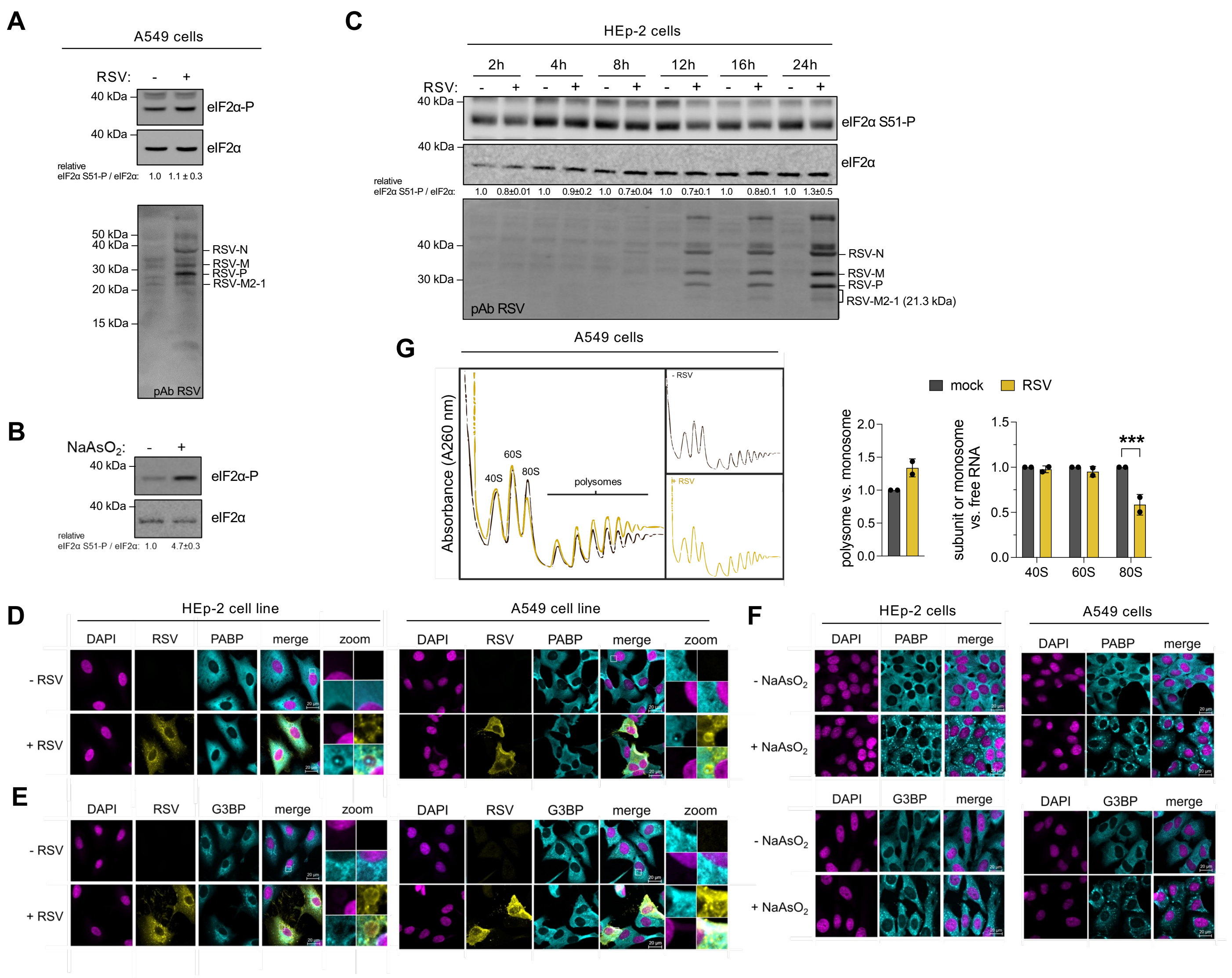

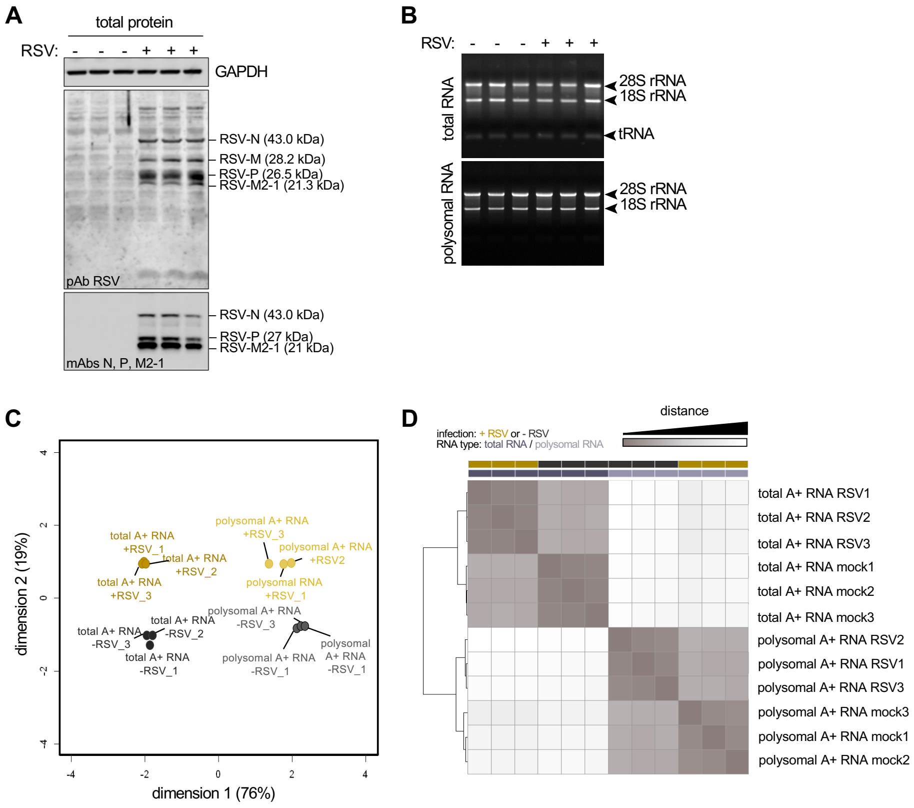

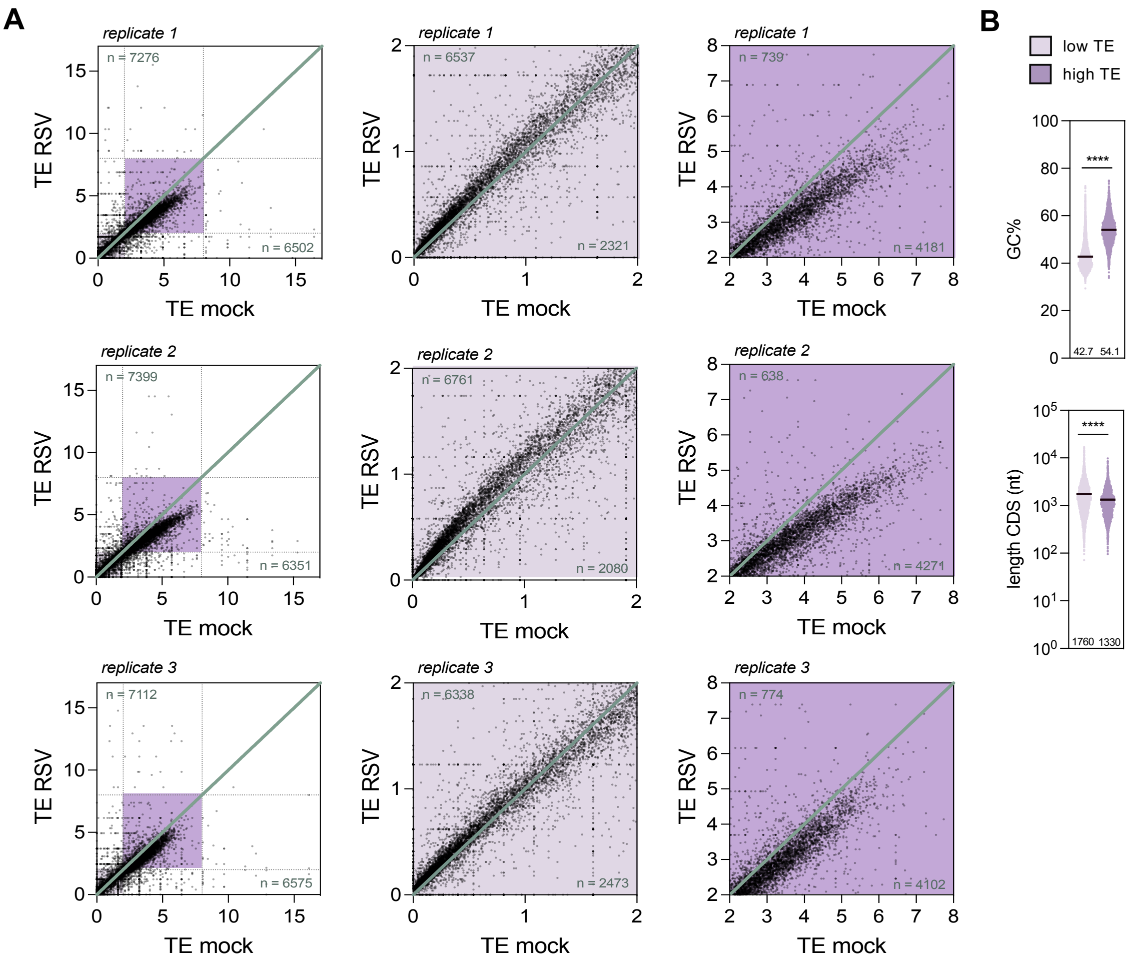

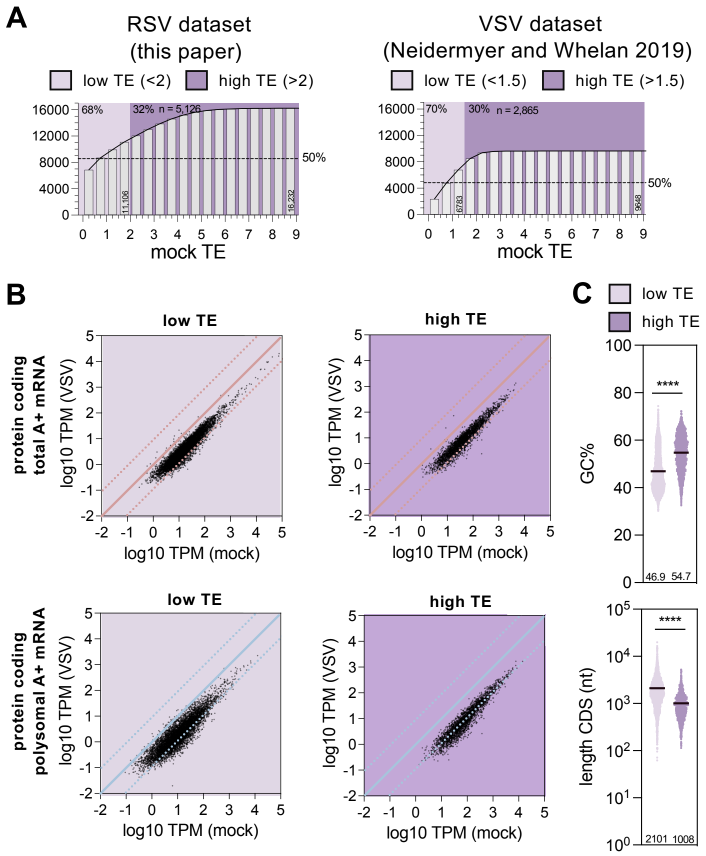

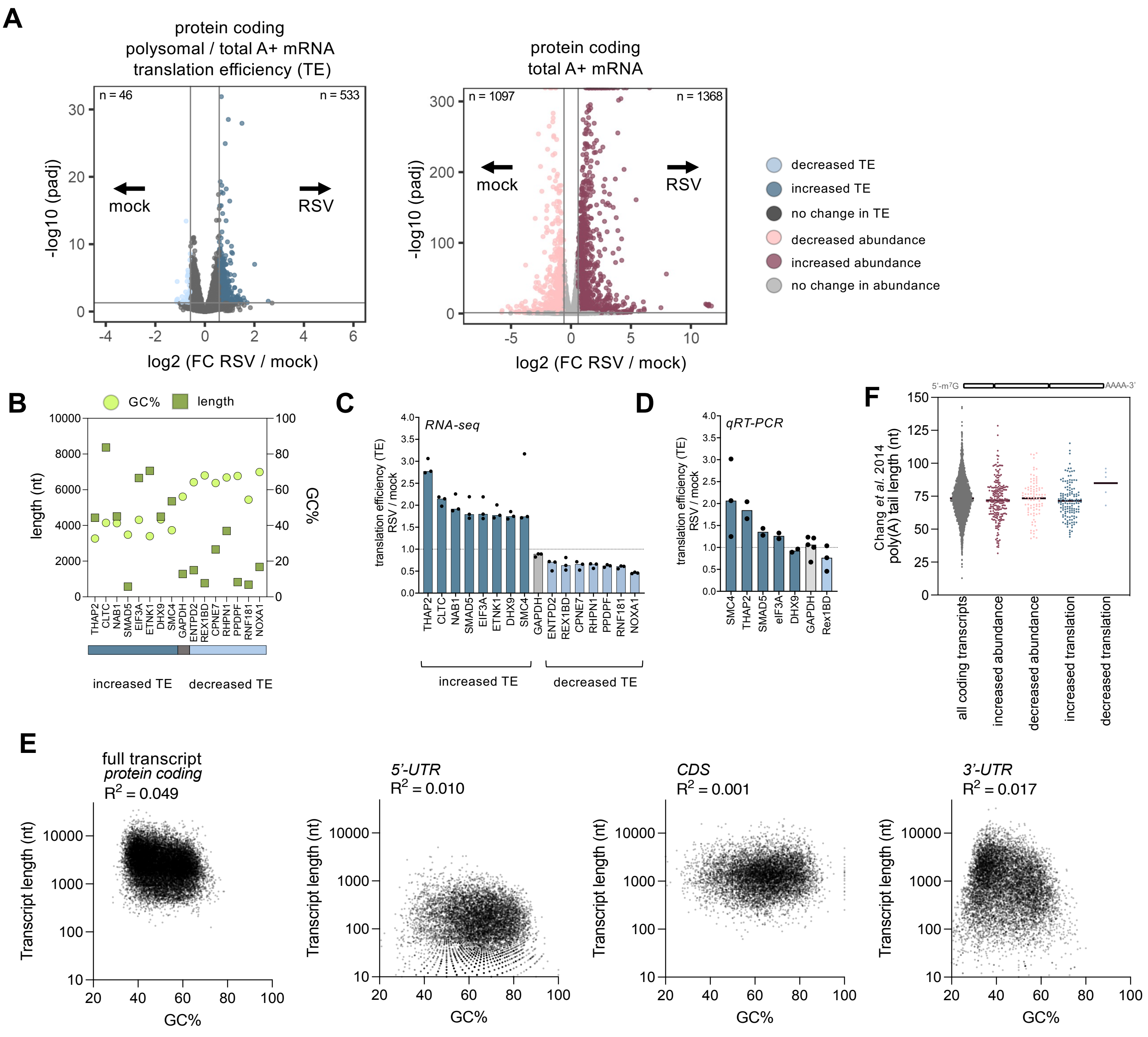

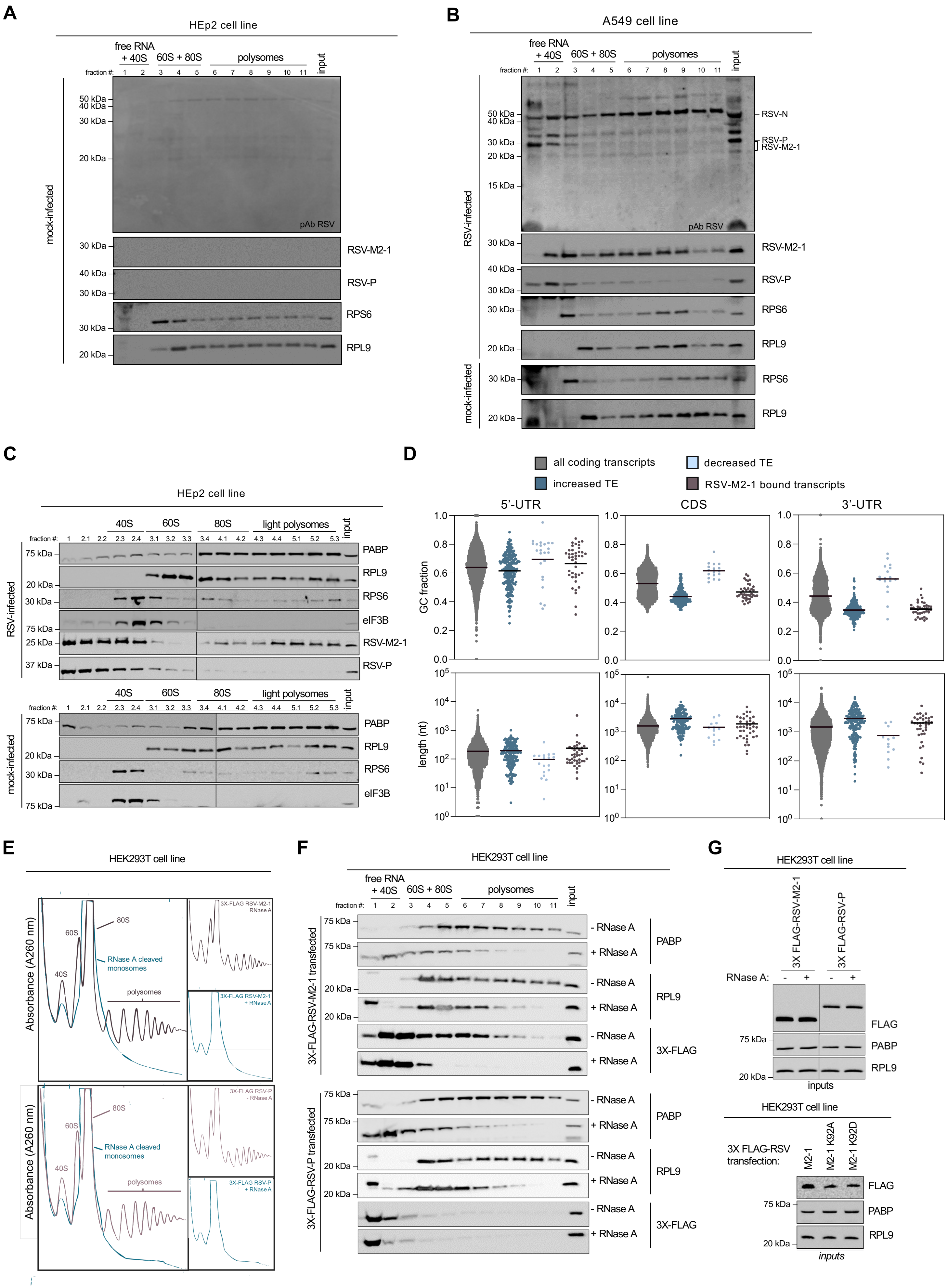
